# Whole-Genome Population Genomics Reveals Lineage Structure and Adaptive Potential of *Philaenus spumarius*, the Principal Vector of *Xylella fastidiosa* in Europe

**DOI:** 10.64898/2025.12.12.693891

**Authors:** Roberto Biello, Sam T. Mugford, Thomas C. Mathers, Qun Liu, The Spittlebug Consortium, Saskia A. Hogenhout

## Abstract

*Philaenus spumarius* (L.), the meadow spittlebug, is the principal European vector of *Xylella fastidiosa*. This xylem-feeding insect has a broad host range, ecological plasticity, and mobility, making it an efficient vector across diverse landscapes. Yet, major gaps remain in understanding its genetic diversity, migration patterns, and local adaptation, limiting effective control of *X. fastidiosa* outbreaks. To address these gaps, we worked with a global network of researchers and citizen scientists to assemble a geographically and ecologically diverse collection of *P. spumarius* samples. We generated a chromosome-level genome assembly for *P. spumarius* and high-quality assemblies for four related spittlebug species, providing a robust genomic framework for evolutionary and epidemiological studies. Resequencing 430 *P. spumarius* individuals from across the globe uncovered boreal, temperate, and semi-arid lineages shaped by geography and climate. The lineages vary in reproductive isolation and in the degree of mito-nuclear divergence. Iberian populations grouped with an individual identified as *P. tesselatus* contributing to regional genetic complexity. Genomic scans identified loci under selection in Apulian populations, including signatures in sulfotransferase (SULT) genes potentially linked to behavioural plasticity, host plant specialisation, or insecticide resistance in olive-growing regions. Migration analyses indicated limited long-distance dispersal but strong local connectivity, consistent with the rapid regional spread of *X. fastidiosa*. Together, these findings reveal the lineage structure and adaptive potential of *P. spumarius*, the key vector of *X. fastidiosa* in Europe. They underscore the importance of targeted surveillance of locally adapted populations and provide genomic tools for monitoring vector dynamics and mitigating emerging disease risks.

## INTRODUCTION

*Philaenus spumarius* (L.) (Hemiptera: Aphrophoridae), commonly known as the meadow spittlebug, is a widespread and highly adaptable xylem-sap–feeding insect that has been extensively studied for its remarkable variation in colour morphs (Stewart and Lees, 2008). Relatively recently, this insect has been recognized as the primary European vector of the plant pathogen *Xylella fastidiosa* (Saponari et al., 2014). Its capacity to feed on over 600 plant genera (Thompson et al., 2023), coupled with its mobility and ecological plasticity, make it a particularly efficient vector of *X. fastidiosa* across both natural and agricultural landscapes (Cornara et al., 2018). Although the biology, life cycle, and behaviour of *P. spumarius* have been well studied, and genomic resources have begun to shed light on its population structure (Rodrigues et al., 2014; Seabra et al., 2021), critical questions regarding its genetic diversity at both local and global scales, migration dynamics, and the ability to adapt to specific plant hosts and environments have remained (Kottelenberg et al., 2021; Bodino et al., 2023; Gillioli et al., 2024). Moreover, genomics and population data for spittlebugs, and sap-feeding insects of the order Hemiptera more generally, remain limited. These gaps hinder the development of precise, sustainable control strategies to limit the impact of *X. fastidiosa* outbreaks.

*X. fastidiosa* is a xylem-limited bacterial pathogen native to the Americas, where it has caused significant economic losses in crops such as citrus and grapevine, known as Citrus Variegated Chlorosis (CVC) and grapevine Pierce’s disease (PD), respectively (Park et al., 2011; Perring et al., 2001). *X. fastidiosa* is transmitted by xylem-feeding insects, and in the Americas, these are primarily sharpshooters (Redak et al., 2004), although *P. spumarius* is also capable of vectoring *X. fastidiosa* causing PD in California (Beal et al., 2021). The highly migratory and polyphagous glassy-winged sharpshooter (GWSS; *Homalodisca vitripennis*) has played a key role in its spread among grapevines in the US, leading to devastating outbreaks in Florida and California (Sisterson et al., 2020; Daugherty and Almeida, 2019). Similarly, the highly polyphagous and abundant *P. spumarius* has played a key role in spreading *X. fastidiosa* in Europe.

The emergence of Olive Quick Decline Syndrome (OQDS) in the Salento Peninsula of southern Italy marked the beginning of one of the most severe plant disease outbreaks in Europe. First observed in 2010 as a mysterious dieback of olive trees, the disease was officially linked to *X. fastidiosa* in 2013, by which time approximately 8,000 hectares of olive orchards were affected, and within twelve months, the infected area had nearly tripled (Saponari et al., 2013; Boscia, 2014; EFSA, 2013; Stokstad, 2015). Likely introduced around 2008 (Kottelenberg et al., 2021), *X. fastidiosa*, identified as subspecies *pauca* strain ST53, spread rapidly, devastating olive orchards across the Apulia region and leading to the death of hundreds of thousands of trees (Bodino et al., 2023; Gillioli et al., 2024). The ability of *P. spumarius* to colonize the olive canopy during dry months, combined with its high population densities and capacity for both active and passive dispersal, has driven rapid disease expansion with major economic and ecological consequences (Cornara et al., 2017; Cornara et al., 2018).

Given its abundance and broad host range, *P. spumarius* is likely the primary driver of *X. fastidiosa* subspecies spread following their introduction into Europe. In 2016, the subspecies *multiplex*, *fastidiosa*, and *pauca* were detected across the Balearic Islands (Olmo et al., 2017), where *P. spumarius* is regarded as the main vector of almond leaf scorch disease (ALSD) and Pierce’s disease (PD). Greenhouse studies have further confirmed its capacity to transmit *X. fastidiosa* between grapevines and almonds (Moralejo et al., 2020). The following year, *X. fastidiosa* subsp. *multiplex* was identified in almond trees in eastern Spain, with subsequent detections in Madrid and in ornamental plants in Alicante (Marco-Noales et al., 2021). While Italy remains the most severely affected region, these scattered findings of diverse genotypes, including *fastidiosa*, *pauca*, *multiplex*, and *sandyi*, underscore the complex, multi-origin nature of *X. fastidiosa* introductions and spread in Europe (Landa et al., 2020; EFSA et al., 2022). Additional outbreaks, generally linked to the trade of infected nursery plants, have been reported in France and Switzerland (2015), Germany (2014; subsequently eradicated), and Portugal (2019) (Auricoste et al., 2017; Denance et al., 2017; Soubeyrand et al., 2018; EFSA, 2015; EPPO, 2019). Given its widespread presence in some of these countries, *P. spumarius* is well positioned to drive further spread of the bacterium.

*P. spumarius* is a key representative of the Cercopoidea superfamily, which comprises around 3000 phytophagous xylem sap-feeding species (Cryan and Svenson, 2010; Crispolon et al., 2023). These insects are widely recognized for producing frothy secretions known as "cuckoo spit," named for their seasonal appearance coinciding with the arrival of cuckoo birds in Europe (Svanberg, 2017). The superfamily Cercopoidea is split into 5 families, including Aphrophoridae and Cercopidae (Cryan and Svenson, 2010). Within Aphrophoridae, other notable genera include *Neophilaenus* (e.g., *N. campestris*) and Aphrophora (e.g., *A. alni*). The family Cercopidae is the largest in the Cercopidae and includes *Cercopis vulnerata* (Crispolon et al., 2023). *Philaenus* species are classified into the “spumarius” and “signatus” groups based on male genitalia (Drosopoulos and Remane, 2000). The “spumarius” group includes *P. spumarius*, *P. tesselatus*, *P. loukasi* and *P. arslani* and the “signatus” group *P. maghresignus*, *P. italosignus*, *P. signatus* and *P. tarifa* (Drosopoulos and Remane, 2000).

*P. spumarius* and *P. tesselatus* thrive on a wide range of plants, especially dicots, whereas others have more specific host preferences. *P. spumarius* is widely distributed across Europe and has been introduced to regions such as the Azores, USA, and New Zealand (Stewart and Lees, 1996; Drosopoulos and Remane, 2000; Yurtsever, 2000; Seabra et al., 2021). *P. tesselatus*, initially described as a distinct species but later as a subspecies with *P. spumarius*, has been reported in Tunisia, Morocco, Algeria, and parts of Southern Europe (Maryanska-Nadachowska et al., 2010; Maryanska-Nadachowska et al., 2012). *P. spumarius* and *P. tesselatus* may be distinguished based on male genitalia morphology, but not at the mitochondrial cytochrome c oxidase subunit I (COI) gene locus, rendering DNA barcoding with this gene ineffective (Seabra et al., 2021). Additionally, evidence of admixture between *P. tesselatus* from Morocco and *P. spumarius* from the Iberian Peninsula suggests ongoing gene flow between the two taxa (Seabra et al., 2021). In North Africa, *P. tesselatus* was found to be most abundant, followed by *N. campestris*, *N. lineatus* and *P. maghresignus*. In Apulian olive orchards, *P. spumarius* is considered the primary vector of *X. fastidiosa*, and other xylem feeders like *P. italosignus*, *N. campestris* and *Cercopis sanguinolenta* have also been recorded, though their low abundance on olive trees and limited transmission efficiency suggests a minor role in the spread of *X. fastidiosa* (Elbeaino et al., 2014; Cornara et al., 2017; Cornara et al., 2020; Cavalieri et al., 2019; Bodino et al., 2019).

As a univoltine species, *P. spumarius* completes one generation annually, overwintering as eggs laid from late August through autumn by polyandrous females, with reported fecundity ranging from 10 to 113 eggs (Mundinger, 1946; Wiegert, 1964; Di Serio et al., 2019). After hatching, nymphs pass through five instars, producing protective spittle masses that shield them from predators and desiccation. These nymphs are capable of limited movement between host plants (Weaver and King, 1954; Bodino et al., 2021), and while peak field densities may reach up to 1280 individuals/m² (Wiegert, 1964), lower densities are more typical across Europe (Zajac and Wilson, 1984). Adult emergence begins in spring and individuals may persist into autumn, with occasional overwintering under mild conditions. Newly emerged adults are pale, gradually developing their characteristic coloration, which varies widely in dorsal pattern and is genetically controlled by major genomic loci and linked to physiological and life-history traits (Rodrigues et al., 2016). Sexual dimorphism in phenology is evident, with males emerging earlier and declining later in the season (Edwards, 1935; Halkka, 1964). Although *P. spumarius* adults are capable of flight, field studies show that their effective dispersal is generally limited. They primarily move by jumping or crawling using powerful hind legs (Burrows, 2003). Early observations reported occasional short flights of up to ∼100 m (Weaver and King, 1954) and activity typically occurring 15–70 cm above ground, with rare movements reaching several metres (Wilson and Shade, 1967). However, recent mark-release-recapture experiments indicate much more restricted natural dispersal. Bodino et al. (2021) found that median displacement after one day was only 26 m in olive groves and 35 m in meadows, and modelling predicted that 50% of individuals remain within 200 m of the release point (98% within 400 m) over a two-month period. Similarly, Casarin et al. (2023) observed that *P. spumarius* mainly moved by short jumps of approximately 1 m, with a maximum recorded displacement of 32 m over 27 days. Adults acquire *X. fastidiosa* soon after emergence and can transmit the bacterium throughout their lives, particularly contributing to secondary spread in olive groves during summer and autumn. Passive dispersal, including via wind and vehicles, and seasonal vertical migration from ground vegetation to canopy-level habitats also play roles in their distribution and epidemiological significance (Weaver and King, 1954). Passive dispersal of the vector has contributed to the fast spread of the OQDS in the Apulia Region of Italy, where the estimated rate of movement of the disease front was 10.0 km per year (Kottelenberg et al., 2021) However little data exist on the medium to longer-term patterns of movement and migration patterns of *Philaenus* that are likely drivers of the spread of *Xylella*.

To address critical gaps in understanding the genetic diversity, migration, and adaptive potential of *P. spumarius*, we generated a chromosome-level genome assembly alongside assemblies for close relatives and analysed resequencing data from 430 individuals worldwide. Our results reveal distinct genetic lineages shaped by geography and climate, high gene flow in southern Italy consistent with rapid *X. fastidiosa* spread, and loci under selection linked to behavioural plasticity and detoxification. These findings provide a genomic framework for assessing the risk of *X. fastidiosa* spread and for developing targeted management strategies.

## RESULTS

### A high-quality chromosome-level genome for the meadow spittlebug *P. spumarius*

To enhance the accuracy of *P. spumarius* population genomic and local adaptation studies, we generated a chromosome-level genome assembly for this species. Illumina, PacBio, 10 X Genomics data from a single individual from Portugal (Fontanelas, Sintra; Biello et al., 2020) and *in vivo* chromatin conformation capture (Hi-C) data from a pooled sample of individuals from the Fontanelas population were used to categorize and order the genome into twelve chromosome scale scaffolds, corresponding to the haploid chromosome number of this species (Kuznetsova et al., 2003) (**Figure 1a**). This resulted in a final assembly of 2.6 Gb, with 92% of assembled content contained in the twelve scaffolds, and a scaffold N50 of 195 Mb (**Table 1**; **Figure 1b**). The lengths of the twelve chromosomes ranged from 431 to 112 Mb. The completeness of the gene space in the assembled genome was assessed using the Benchmarking Universal Single Copy Orthologues (BUSCO) pipeline (Waterhouse et al., 2018), with 97.1% of the Hemiptera gene set found to be present as complete single copies (**Table 1**; **Figure 1b**). Thus, the new assembly (Phspu_JIC_v2) represents a near complete, highly contiguous assembly and a significant improvement on the existing assembly of this species (Phspu_JIC_v1; Biello et al., 2020). K-mer analysis was employed to evaluate the accuracy and completeness of the genome assembly (**Figure S1)**. The quality value (QV) of the genome was 32.25 (accuracy of 99.94% or an average of one error per 1680 bases). Blobtools analyses revealed scaffolds mostly from Hemiptera order, confirming the absence of contaminants in the assembled chromosomes (**Figure S2**). These assessment results further validated the completeness of the *P. spumarius* genome assembly.

**Figure 1.**
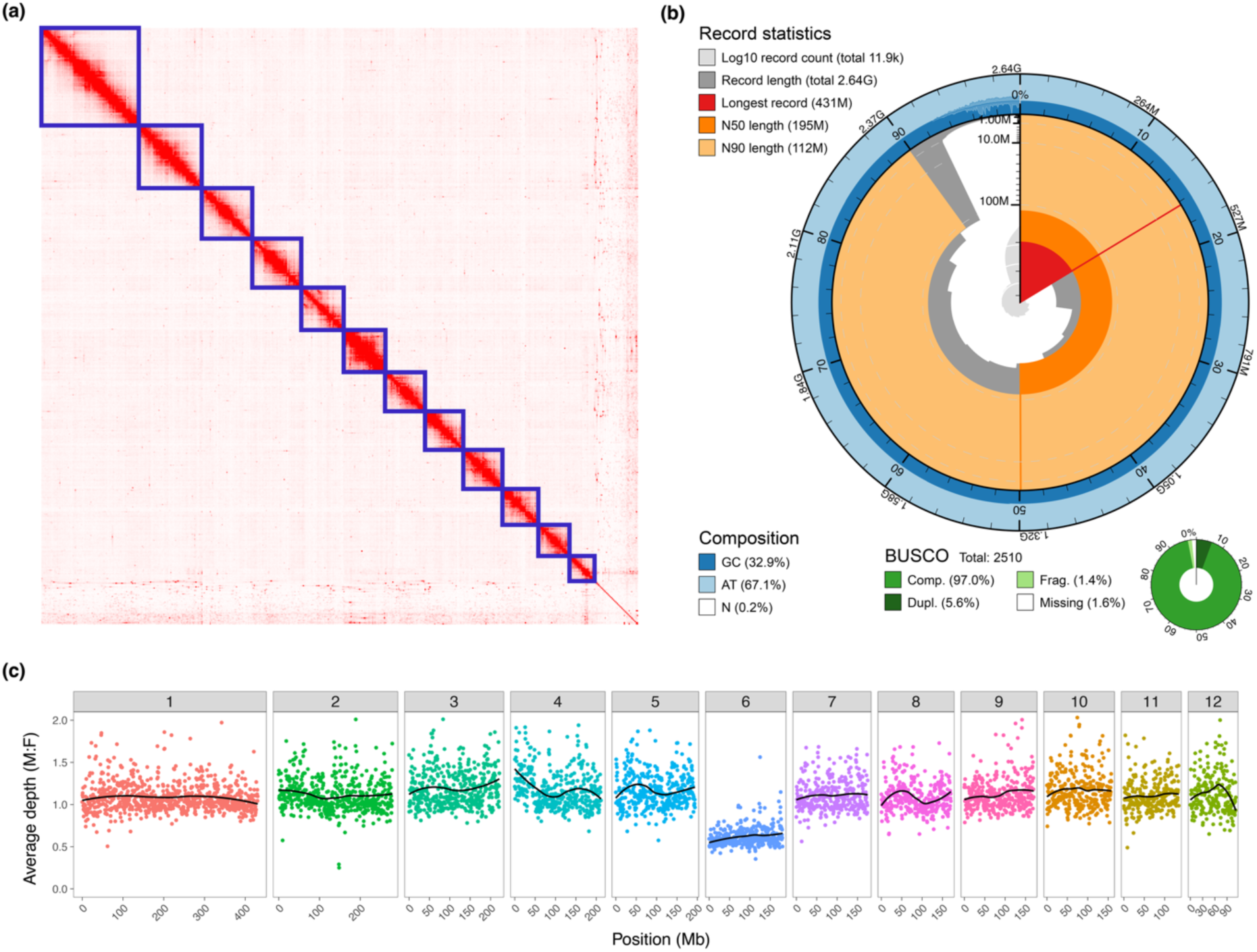
Chromosome-level genome assembly of *Philaenus spumarius* (Phspu_JIC_v2). (a) Heatmap showing the frequency of Hi-C contacts along the Phspu_JIC_v2 genome assembly. Blue lines indicate super scaffolds and green lines show contigs. Genome scaffolds are ordered from longest to shortest with the x- and y-axis showing cumulative length in millions of base pairs (Mb). (b) BlobToolKit Snail plot showing a graphical representation of the quality metrics presented in **Table 1** for Phspu_JIC_v2 assembly. The longest scaffold (red vertical line), N50 (orange track), N90 (light orange track), GC content (external blue track) and BUSCO scores (outer circular pie chart in green) are showed. (c) Identification of sex chromosomes in *P.spumarius*, with scaffold 6 assigned as the sex chromosome based on the mapping of genomic resequencing reads from three individuals of each sex to the reference genome. The y-axis shows the ratios of male-to-female read coverages across the 12 scaffolds.

**Table 1.** Genome assembly statistics of five spittlebug species.

|  | *Philaenus*<br>*spumarius* | *Philaenus*<br>*tesselatus* | *Philaenus*<br>*maghresignus* | *Aphrophora*<br>*alni* | *Cercopis*<br>*vulnerata* |
| --- | --- | --- | --- | --- | --- |
| Base pairs (Mb) | 2,635 | 2,800 | 2,741 | 1,690 | 750 |
| Number of scaffolds | 11,927 | 12,870 | 14,790 | 4,297 | 39,131 |
| Scaffold N50 (Mb) | 194 | 0.720 | 0.553 | 133 | 0.297 |
| Scaffold L50 | 5 | 1,063 | 1,339 | 6 | 609 |
| Longest scaffold (Mb) | 431 | 5.1 | 4.9 | 167 | 2.9 |
| Complete BUSCOs<br>(hemiptera_odb10) | 2,434 (97%) | 2,435 (97%) | 2,442 (97.3%) | 2,420<br>(96.4%) | 2,318<br>(92.3%) |
| Complete and single-copy BUSCOs | 2,293 | 2,296 | 2,375 | 2,363 | 2,280 |
| Complete and<br>duplicated BUSCOs | 141 | 139 | 67 | 57 | 38 |
| Fragmented BUSCOs | 36 | 45 | 40 | 23 | 100 |
| Missing assembly<br>BUSCOs (M) | 40 | 30 | 28 | 67 | 92 |

Read coverage depth of female and male individuals identified scaffold 6 as the *P. spumarius* sex chromosome based 50% lower read coverage compared to the autosomes in males (**Figure 1c**), in agreement with the XX (female)/X0 (male) sex determination of *P. spumarius* (Felt et al., 2017).

The *P. spumarius* genome has a G + C content of 32.9%, which is similar to the other Cercopoidea species (Chen et al., 2022). The circular mitochondrial genome was 15,310 bp in length, with 38 genes typical of insect mitochondrial genomes, including 13 protein coding genes, two ribosomal RNAs (rRNA), 23 transfer RNAs (tRNA) and an A + T rich region (**Figure S3**).

Structural genome annotation using a workflow incorporating RNAseq and UniProt data predicted a total of 29,776 protein-coding genes in the assembly. Of these, 25,525 were assigned functional annotations (**Table S1**). We also annotated 1,587 Mb of repetitive elements (60.2% of the genome) in this genome (**Table S2**).

To further investigate *P. spumarius* and *P. tesselatus* relationships, we also sequenced the genomes of *P. tesselatus* and *P. maghresignus*. These species were selected based on their overlapping or adjacent geographic locations to *P. spumarius* with *P. tesselatus* found in southern Spain and Portugal and north Africa (Nast, 1972) and *P. maghresignus* in north Africa (Drosopoulos, 2003). The assembly genome sizes of the three species were similar (**Table 1**; see **Figure S4-S6** for details on whole- and mitochondrial genome assemblies) and they showed comparable levels of protein annotated and repetitive element content (**Table S1-S2**).

### Genomic and phylogenetic insights into spittlebug evolution

To place these new genome assemblies in a phylogenetic context and to investigate gene family evolution across the Auchenorrhyncha, we compared the proteomes of *Philaenus* spp. (comprising the complete set of annotated protein-coding genes) to those of 14 other Hemiptera species with fully sequenced genomes (**Table S3**). To comprehensively investigate the relationships between *Philaenus* species and other Auchenorrhyncha species, we sequenced the genome of a member of the Cercopidae family, *Cercopis vulnerata*, and a representative of the Aphrophoridae family, *Aphrophora alni* (**Table 1**; see **Figure S4-S6** and **Table S1-S2** for details on genome assemblies and annotations). We also included the genome sequence of the rice spittlebug, *Callitettix versicolor* (Chen et al., 2022). Since the annotation for the *C. versicolor* genome was not publicly available, we provided the annotation herein (see **Table S1-S2**). This extended dataset allows for a more robust exploration of the genetic relationships within the Auchenorrhyncha suborder, enhancing the depth of our analysis.

Our maximum likelihood phylogenetic analysis, based on a concatenated alignment of conserved single-copy genes, produced a species tree with most of nodes with 100% bootstrap support except those indicated by asterisks (support values: 78–98%) (**Figure 2a**). In total, we clustered 236,619 proteins into 22,454 orthogroups (gene families) and 11,539 singleton genes (**Table S4**) within the Auchenorrhyncha suborder. Among the 29,776 predicted genes in the *P. spumarius* genome, 14,477 (64.5%) have an orthologue in at least one other Auchenorrhyncha species, and 201 species-specific orthogroups (**Table S5**). Hemiptera genomes are known to be subject to high levels of ongoing gene duplication (Fernández et al., 2020; Julca et al., 2020; Mathers et al., 2017; Thorpe et al., 2018; Biello et al. 2021). *Philaenus* spp. are no exception, as we identified a significant number of lineage-specific gene families within *Philaenus* genus and a range of 3,804-6,268 genes that have undergone lineage-specific duplications (**Table S6**).

**Figure 2.**
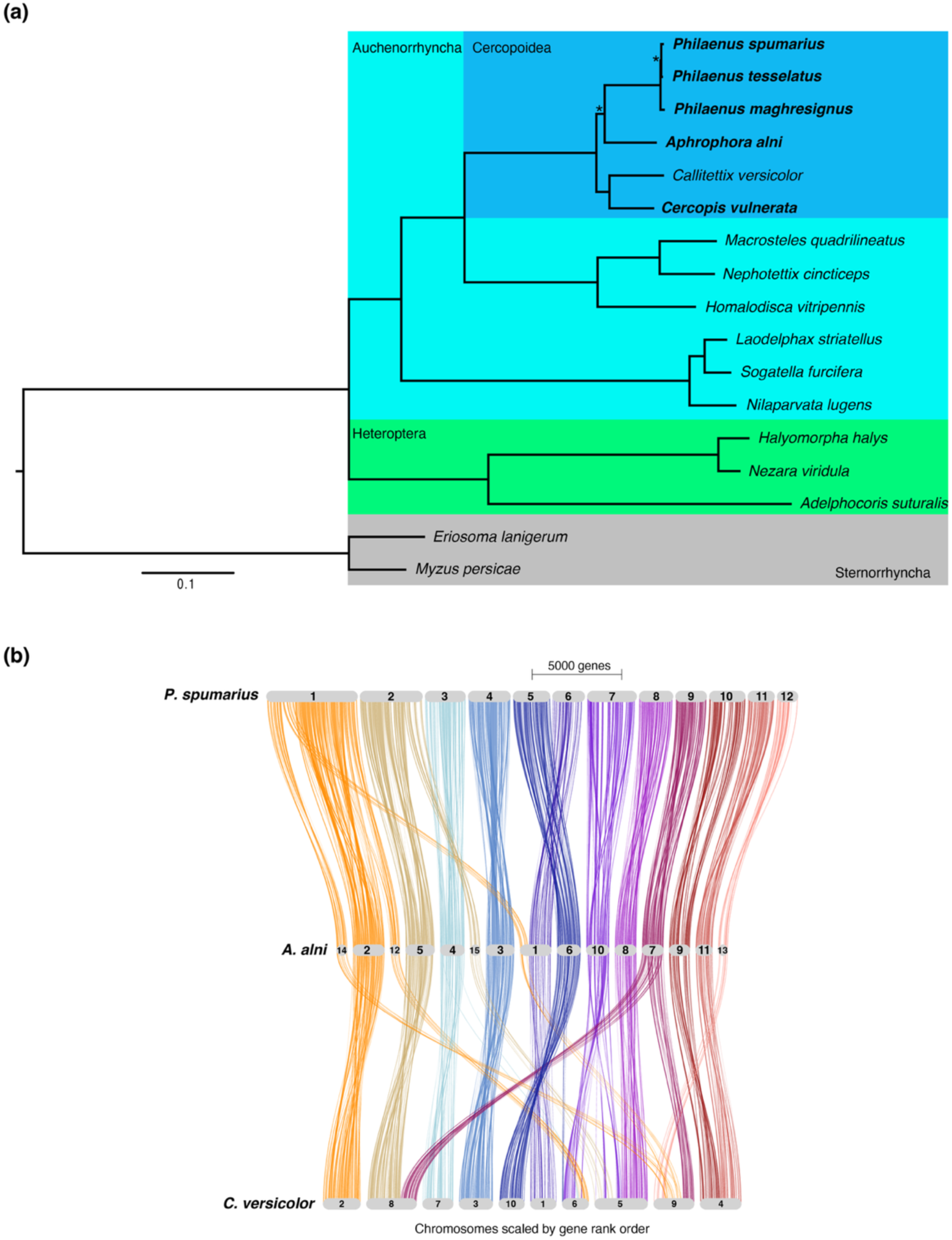
Phylogeny and genome reorganization. (a) Maximum likelihood phylogeny of *Philaenus spumarius*, *P. tesselatus*, *P. maghresignus*, *Aphrophora alni*, *Cercopis vulnerata* (in bold) and 12 other Hemiptera species based on a concatenated alignment of 93 conserved one-to-one orthologues. The tree is rooted with the aphid branch. Clades are coloured by taxonomic groups. Branch lengths are in amino acid substitutions per site. All nodes are supported by 100% bootstrap values, except those marked with asterisks, which have support values ranging from 78% to 98%. (b) Genome reorganization across Cercopoidea species. Pairwise synteny relationships are shown between *P. spumarius* and the chromosome-scale genome assemblies of *A. alni* and *Callitettix versicolor*.

The *P. spumarius* and *A. alni* chromosomes are highly collinear and largely syntenic, with little evidence of major rearrangements (**Figure 2b**). Ancestral chromosomal fission likely contributed to the increased chromosome number in *A. alni* relative to other Cercopoidea. For instance, PsChr1 corresponds to AaChr2, AaChr12, and AaChr14, while PsChr2 aligns with AaChr5 and AaChr15 (**Figure 2b**). On average, chromosomes are longer in *P. spumarius* than in *A. alni* or *C. versicolor*, consistent with the larger *P. spumarius* genome. By contrast, *C. versicolor* exhibits a more rearranged karyotype, reflecting its more distant evolutionary relationship as a member of the Cercopidae, whereas *P. spumarius* and *A. alni* both belong to the Aphrophoridae.

### Global genetic diversity in *P. spumarius* through resequencing of 430 samples

We engaged with a global network of colleagues and citizen scientists to curate a diverse collection of *P. spumarius* samples from various locations worldwide. Participation opportunities were promoted through online seminars, workshops, and social media channels. Participants received kits containing collection tubes and detailed instructions for sample collection, including guidelines for precisely recording collection locations (Wouters et al., 2020). Upon receiving the returned collection tubes, each containing a single desiccated individual, meticulous records were maintained. The insects were then snap-frozen in liquid nitrogen and stored at -80°C to ensure their preservation.

A significant proportion of the received samples originated from *P. spumarius* nymph specimens collected from lavender in gardens. However, we also received samples from *P. spumarius* nymphs collected from diverse plant hosts, as well as samples representing a variety of spittlebug species. Therefore, the collection represents a comprehensive representation of the species across different environments and hosts, enriching the diversity and depth of our study.

From the collected samples, 434 individuals were processed for DNA extraction and submitted for genome resequencing at ∼4 individuals per location (**Figure S7, Table S8**). Each genome was sequenced using the Illumina sequencing platform, generating 150-bp paired-end raw reads. A total of ∼30 Gb of quality-filtered reads for each individual were aligned to the *P. spumarius* reference genome (Phspu_JIC_v2). Individuals with low mapping rates or insufficient coverage (<75%) were excluded, resulting in 430 individuals retained for variant discovery.

After further filtering, we identified 9,600,824 high-quality biallelic single-nucleotide polymorphisms (SNPs). The density of SNPs was similar across the twelve chromosomes averaged between 3,004 and 1,364 SNPs every 100 kb of genomic length, except the sexual chromosome (scaffold_6) that averaged around 151 SNPs every 100 kb (**Figure S8**). To explore the genetic relationships among individuals and confirm the identification of our samples as *P. spumarius*, we first constructed a phylogenetic tree based on its approximately 15.3 kb whole-genome mitochondrial sequence (**Figure S3**). Across 430 individuals, the median of the per-individual mean sequencing depth for the mitochondrial genome was 978× (**Figure S9**). We identified three primary lineages, largely matching the defined Eastern Mediterranean, Western, and Northeastern haplogroups (Rodrigues et al., 2014; Seabra et al., 2021) (**Figure S10**). The sample identified as *P. tesselatus* clustered among the *P. spumarius* samples from Spain, France, Sardinia and Corsica (**Figure S10**), in agreement with mitochondrial genotypes not distinguishing *P. spumarius* and *P. tesselatus* individuals (Seabra et al., 2021).

Principal Component Analysis (PCA) of nuclear genome-wide polymorphisms revealed distinct clustering patterns among the studied populations. The first principal component (explaining 23.8% of the genetic variance) identified *P. tesselatus* and individuals from Spain as a cohesive group, which clustered separately from other populations. Portuguese samples bridged the two clusters (**Figure 3a**), suggesting that individuals in Spain represent *P. tesselatus* and that admixture has occurred between the *P. tesselatus*-like populations in southern Spain and *P. spumarius* in Portugal. Exclusion of the *P. tesselatus*-like individuals from Spain revealed a second principal component of an east-to-west gradient of genetic relatedness across Europe, partially separating populations from Turkey, Greece, and Croatia (**Figure 3a, Figure S11**).

**Figure 3.**
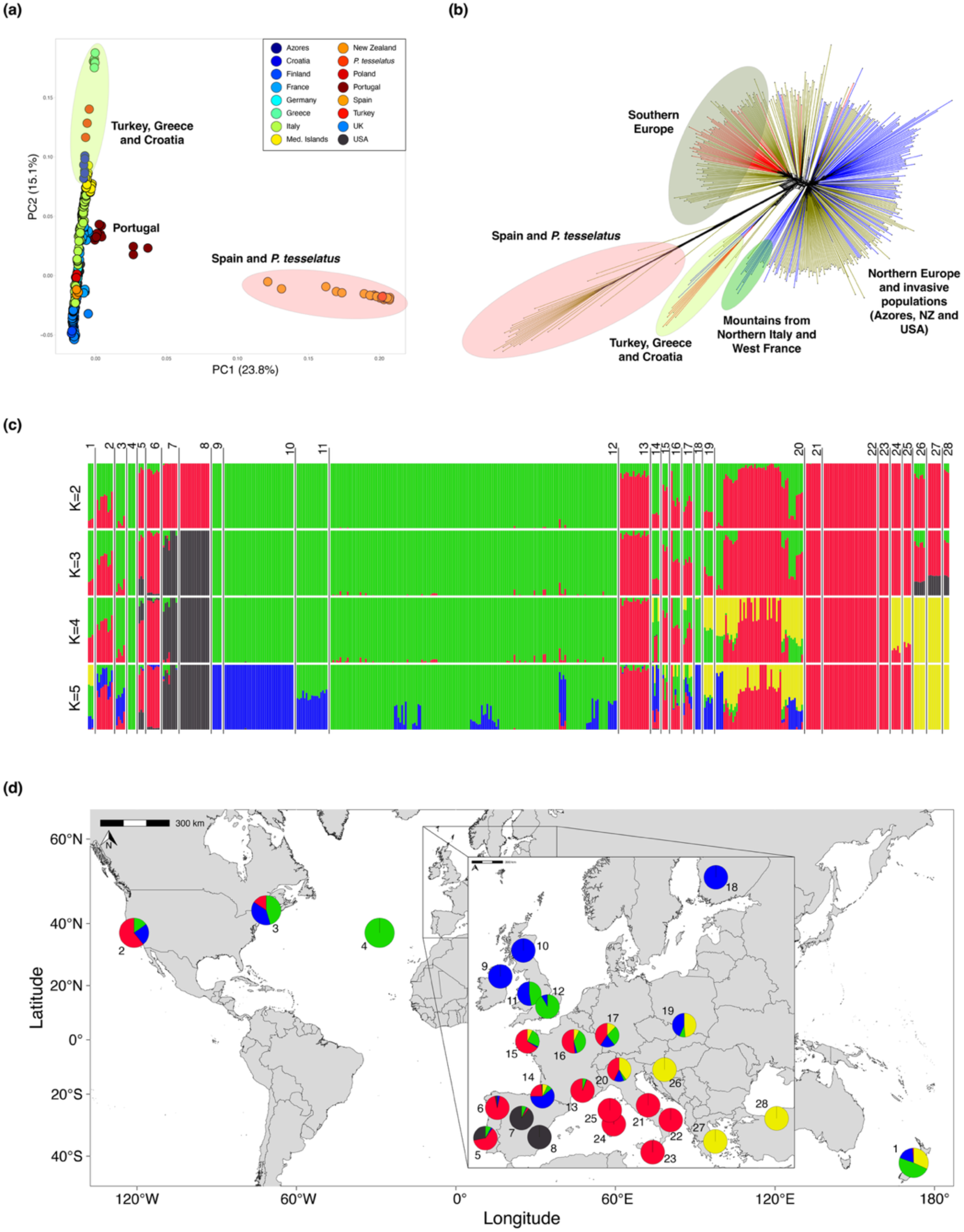
Population structure of *P. spumarius*. (a) PCA based on the full data set showing the first two PCs. Individuals are represented as dots, coloured by geographical origin reflecting the legend. (b) Phylogenetic network inferred using the Neighbour-Net method based on genome-wide SNPs. Branches are coloured following the mitochondrial haplogroups (see **Figure S10**). (c) Individual genetic assignment as inferred by structure from K = 2 to K = 5. Bar plot is divided in 28 sampling sites, with nearby locations clustered into a single site, showed in the map. (d) Geographic distribution of 28 sampling sites (circles). The pies indicate the ancestral components inferred using ADMIXTURE when K = 5.

Phylogenetic network analysis based on nuclear genome-wide polymorphisms further supported these findings. Samples from southern Spain and the *P. tesselatus* sample grouped together forming a distinct clade, while populations from Turkey, Greece, and Croatia grouped into another clade (**Figure 3b**). Notably, samples from high-elevation regions in northern Italy and southwestern France formed a distinct clade (**Figure 3b**), suggesting these populations may represent ancient lineages that have remained isolated mainly from recent population expansions. This isolation may be due to adaptations to colder conditions. The network analysis also highlighted two major groups: one encompassing Southern Europe and another representing Northern Europe (**Figure 3b**). The Northern Europe group also included invasive populations introduced to the Azores, New Zealand, and the USA.

Analyses of cluster assignment tests using LD-filtered SNPs revealed clear geographic structuring of the data, accompanied by varying degrees of admixture events (**Figure 3c**). The lowest cross-validation error (CVe) was observed at K = 3 (**Figure S12**), where individuals collected in Spain formed a distinct cluster (grey bars, **Figure 3c**) from those elsewhere in the Mediterranean (red bars, **Figure 3c**) and northern Europe (green bars, **Figure 3c**). Individuals in southern Spain exhibited no obvious levels of admixture, while those in central Spain, Portugal, Turkey, Greece, and Croatia displayed moderate to high levels of admixture. However, K = 5 best matched the PCA and phylogenetic network analyses based on genome-wide SNPs. Therefore, we concluded that K = 5 provided the best resolution for defining the global *P. spumarius* population structure (**Figure 3d**), represented by: (i) Southern and central Italy, and Sicily (red circles, **Figure 3d**); (ii) Northern Europe, further split into two subgroups: Scotland, Northern Ireland, and Finland (blue circles, **Figure 3d**), and England (green circles, **Figure 3d**); (iii) Southern Spain (black circle, **Figure 3d**); and (iv) Eastern Europe (yellow circles, **Figure 3d**). Individuals from Portugal, central Europe (including northern Italy), the USA, and New Zealand exhibit varying degrees of admixture among these populations. Populations in England showed admixture with more northern populations (**Figure 3d**) and a detectable, though occasional admixture with populations from southern Europe (**Figure 3c**). Notably, isolated populations at high elevations in the Pyrenees and Alps were primarily associated with the Northern Europe cluster (blue circles, **Figure 3c, d**). The ‘blue’ cluster may represent older lineages that spread across Europe earlier than the other populations and may be better adapted to colder climates.

Samples from the Azores clustered closely with populations from southern England and those from New Zealand and the USA displayed admixed genetic profiles that aligned most closely with Central European and the UK populations (**Figure 3c, d**). These patterns suggest that the invasive population in the Azores likely originated from southern England, potentially through a single colonization event. In contrast, the genetic admixture observed in the USA and New Zealand populations indicates that they may have originated from different introduction events involving diverse European source populations. This pattern is consistent with the complex colonization history of these invasive populations and underscores the role of human-mediated dispersal in shaping their current genetic structure.

To assess genetic distances between populations, we calculated pairwise FST values. These analyses revealed significant genetic differentiation between the *P. tesselatus*-like populations from southern and central Spain and the other populations. Including these two locations, FST values ranged from 0.177 between central Spain and central Portugal to 0.405 between southern Spain and Finland (**Figure S13**). In general, the highest levels of differentiation were observed between southern Spanish and non-Spanish populations, consistent with our data showing that southern Spanish populations are *P. tesselatus*-like individuals. Among non-Spanish populations, FST values were comparatively lower, not exceeding 0.234, with greater genetic differentiation typically observed between populations separated by larger geographic distances (**Figure S13**). These findings suggest a pattern of isolation by distance, where genetic divergence increases with geographic separation.

Populations from southern Spain exhibited the lowest levels of heterozygosity, similarly to those of the *P. tesselatus* sample, indicating a high degree of inbreeding and low admixture levels with other individuals identified as *P. spumarius* (**Figure S14**). In contrast, spittlebug populations from central Spain exhibited a wide range of heterozygosity levels (from relatively high to low), likely reflecting the higher levels of admixture observed in this region. Other populations with low heterozygosity included those from northern areas, such as Finland and Scotland, with the Shetland samples showing particularly low values—likely due to their geographic positions (**Figure S14**). Populations from the Azores, USA, and New Zealand also exhibited reduced heterozygosity compared to mainland European populations. This pattern is consistent with founder effects, which occur when new populations are established by a small number of individuals, leading to reduced genetic diversity. These results support the hypothesis that these populations are the result of recent introductions, likely originating from limited source populations in Europe (**Figure S14**).

### Dispersal dynamics and gene flow barriers of *P. spumarius* across Europe and within Italy and the UK

Although *P. spumarius* has limited flight capacity, it is an exceptional jumper and is frequently observed hitchhiking on vehicles, facilitating its regional dispersal. This behaviour is consistent with the admixed populations observed in central Europe. To further investigate *P. spumarius* dispersal patterns within regions, we utilized Estimating Effective Migration Surfaces (EEMS) and Fast Estimation of Effective Migration Surfaces (fEEMS) to visualize spatially heterogeneous isolation-by-distance on a geographic map (Petkova et al., 2016; Marcus et al., 2021).

At a Europe-wide scale, we identified regions with both high and low migration rates (**Figure 4a**). As expected, areas with high migration rates largely corresponded to the geographic distributions of individuals belonging to similar genetic clusters (**Figure 3a**). In eastern Europe, the Carpathian Mountains emerged as a significant barrier to gene flow, restricting genetic exchange between *P. spumarius* populations in Turkey, Croatia, and Greece and those in northern Europe, such as Poland and Finland (**Figure 4a**).

**Figure 4.**
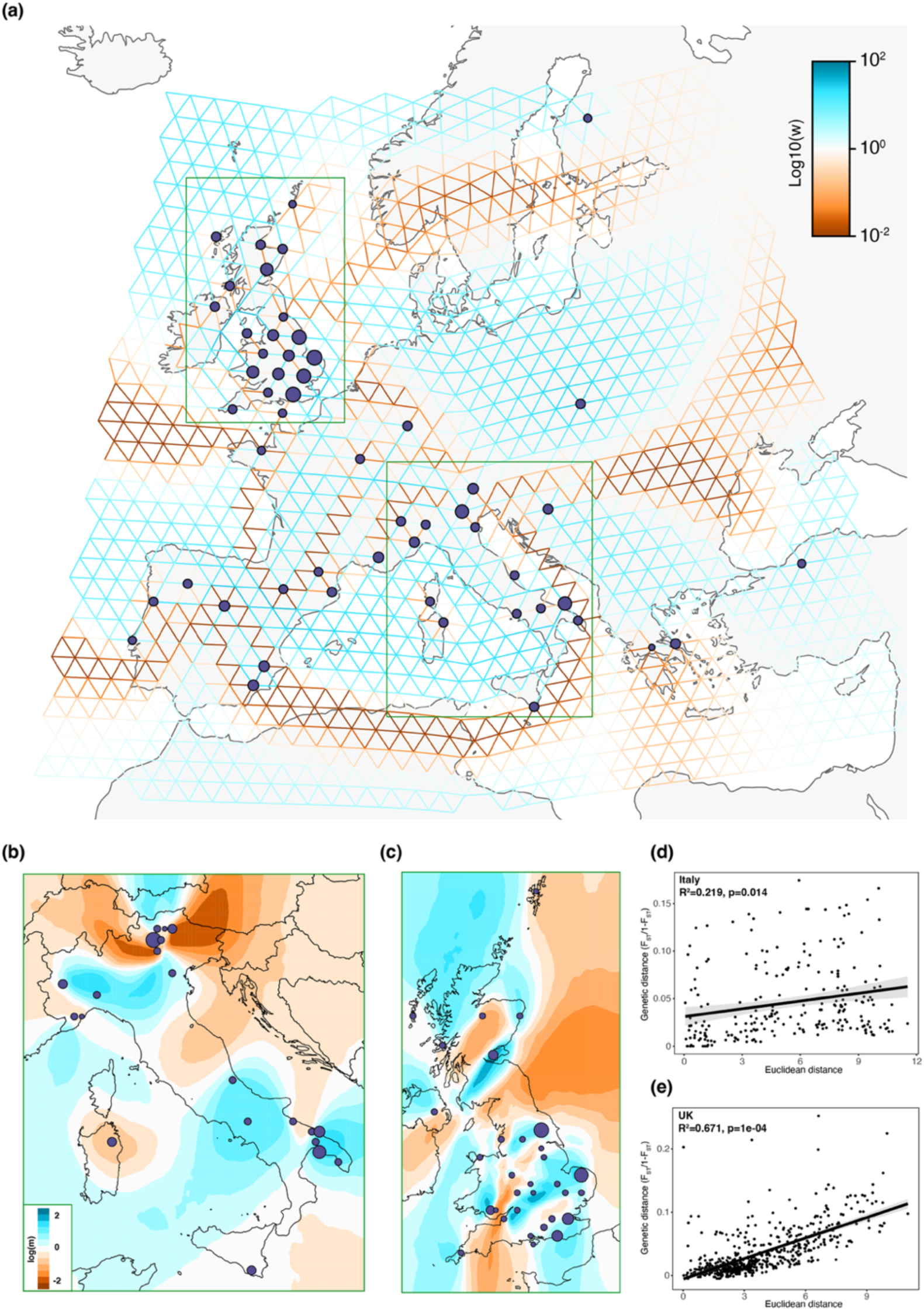
Patterns of migration rates in *P. spumarius* across Europe. Effective migration rates estimated using fEEMS and EEMS, with the fitted parameters displayed on a log scale; lower migration rates are shown in orange, and higher migration rates in blue. (a) Effective migration rates in Europe. (b) Effective migration rates in Italy, and (c) in the UK, presented using the same colour scheme. (d) Regression of pairwise population F_ST_ against geographical distance in Italy, and (e) in the UK.

In contrast, central and western Europe displayed a broader pattern of admixture, although regions of reduced gene flow were notable around the Alps and the Pyrenees (**Figure 4a**). Interestingly, populations in Spain and Portugal exhibited reduced gene flow despite the lack of apparent topographical barriers. This pattern is likely attributable to the prevalence of *P. tesselatus* in southern Spain, which may be influencing genetic structure in these regions.

We further investigated effective migration surfaces within large areas colonized by populations belonging to the same genetic clusters, such as mainland Italy, Sicily, Corsica, and Sardinia (associated with the ‘red’ cluster, **Figure 3d**), and the United Kingdom (associated with the ‘green’ and ‘blue’ clusters, **Figure 3d**). High migration rates and gene flow were detected among populations in the heel of Italy, including Apulia—where the *X. fastidiosa* outbreak in olive trees is prevalent—as well as within central and northern Italy (**Figure 4b**). Apulia, in the heel of Italy, the highest migration rates were observed between the southern Lecce province where *X. fastidiosa* was initially detected and more northern areas of the heel, comprising the Taranto, Brindisi and Bari provinces, consistent with the documented northward spread of *X. fastidiosa* over a five-year period (Bosso et al., 2016; Loconsole et al., 2014; Martelli, 2015). Given that *P. spumarius* is the primary vector for *X. fastidiosa* in olive trees (Saponari et al., 2014; Cornara et al., 2017), this area of highest effective migration likely represents the region traversed by these insects during the same time frame.

Effective migration surfaces indicated that *P. spumarius* populations in southern and central Italy are separated by a barrier in the northeast, possibly because of the highly populated Bari area, while gene flow is higher between these populations and those in the southwest and north, towards the Basilicata, Campania, Molise and Abruzzo regions. This pattern suggests that, historically, *P. spumarius* likely migrated between Apulia and central Italy via the southwestern region towards the north, implying that *X. fastidiosa* may also spread to central Italy through this route. Low migration rates were detected between Apulian and central populations and those in Sicily, Sardinia, and northern Italy. In northern Italy, populations in the northeast and northwest showed limited gene flow, consistent with the northeast populations being collected from mountainous areas at various elevations.

Similar analyses of UK populations revealed regions with both high and low migration rates (**Figure 4c**). Barriers to gene flow were identified in the London area, the higher altitude regions of northern England, between England and Wales, and in Scotland. These barriers align with the distribution of the ‘green’ and ‘blue’ genetic clusters in these regions (**Figure 3b**). Between these barriers, regions with high gene flow were observed, forming migration “corridors” that may facilitate relatively rapid movement of *P. spumarius*, at rates comparable to or higher than those observed in Apulia, Italy.

Isolation-by-Distance (IBD) analysis revealed a stronger positive association between geographic and genetic distance in the UK (Mantel r = 0.671, p = 1e-04) compared to Italy (Mantel r = 0.219, p = 0.014) (**Figure 4d, e**). This suggests that dispersal appears to be more limited across the UK, leading to stronger spatial genetic structure, whereas populations in Italy show weaker spatial genetic differentiation, possibly due to higher connectivity or more frequent gene flow.

### Adaptive evolution of the sulfotransferase genes in Apulia populations

*P. spumarius* is commonly found on olive trees in Apulia, which may reflect its adaptation to the region’s extensive olive monoculture. To identify genes potentially under selection in the Apulian population, we applied the Population Branch Statistic (PBS) approach (Jiang and Assis, 2020; Fabbri et al., 2025) to measure genetic differentiation between the Apulian population and those in Northern Italy, Norfolk, and Spain. We focused on genomic windows with extreme divergence (top 1% PBS values) (**Figure 5a**), identifying 6,791 highly divergent regions. Out of these, we analysed the ones in which genes were annotated, identifying contained 389 protein-coding genes. Functional enrichment analysis (P < 0.05) revealed that these genes were significantly associated with sulphur group transfer activities, including sulfation and sulfotransferase functions (**Figure 5b**).

**Figure 5.**
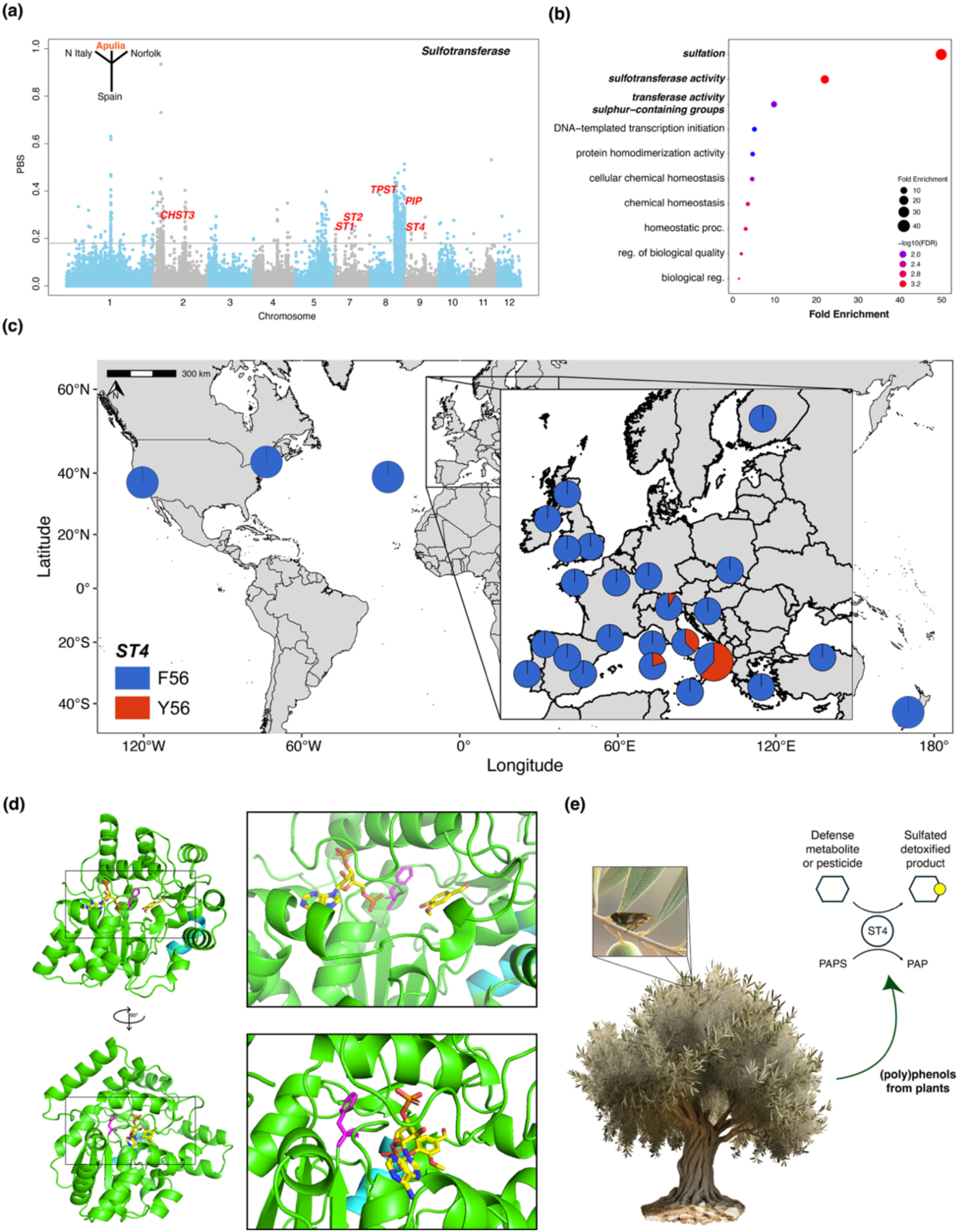
Genomic signatures of selection in the Apulia population. (a) Genomic differences based on population divergence (PBS) among Apulia, Northern Italy, Norfolk and Spain. In bold the outliers gene involved in the sulfotransferase pathway. Statistics were calculated using a 50-kb sliding window. (b) The plot shows the GO categorization of identified genes in significant pathways (FDR < 0.05). The colour of the circle represents the −log (FDR) value. The size of the circle represents the number of selected genes in the enriched pathway. (c) Pie charts of frequency of the nonsynonymous SNP (F56Y) in ST4 gene in different populations. (d) The structural model of the *P. spumarius* ST4 protein (green) in complex with the substrates PAPS and vanillin (yellow) was modelled using Alphafold2 and aligned with the *A. gambiae* vanillin sulfotransferase (7r0u, not shown). The variable positions F56 is coloured in magenta (the Phe sidechain is displayed), and FSQQDGFY 33-42 in cyan (showing the backbone only). The two views show the proximity of the F56 sidechain to the substrate binding pocket, showing the whole protein (left) and a zoomed-in view of the active site (right). (e) Schematic representation of the sulfotransferase pathway in olive. The olive tree and insect were generated using ChatGTP.

Among the genes with the highest PBS values were five of the six annotated sulfotransferases (SULTs) in the *P. spumarius* genome, ST1 through 4, TPST and PIP (**Figure 5a**). SULTs are key enzymes involved in the detoxification and elimination of various endogenous and exogenous molecules (Yamamoto et al 2022). Notably, one of these, sulfotransferase 4 (ST4), exhibited non-synonymous polymorphisms in its protein-coding region. The annotated ST4 gene (PhSpu_g27872.t1) appeared incomplete, missing a conserved C-terminal domain, which was present in an adjacent gene (PhSpu_g27871.t1), suggesting a potential gene model split in the Braker annotation. RT-PCR and Sanger sequencing using primers spanning these two predicted gene models confirmed that they encode a single cDNA corresponding to a full-length ST4. The ST4 coding sequence contained non-synonymous polymorphisms in the 32-FSQQDGFY-42 variable region and a single amino acid polymorphism, F56Y.

Analysis of the prevalence of the F56Y polymorphism across *P. spumarius* populations in Europe revealed that the Y56 variant is more frequent in Apulian populations but occurs at lower frequencies in other regions of Italy. Populations elsewhere in Europe predominantly carry the ancestral F56 variant (**Figure 5c**). The F56Y polymorphism might be an adaptation in the Apulian population to detoxify pesticides or olive-specific defence metabolites.

To better understand the structural implications of these polymorphisms, the ST4 protein of *P. spumarius* was modelled using AlphaFold2 and aligned to the crystal structure of a related sulfotransferase from *Anopheles gambiae* (AgST) in complex with its substrates, vanillin and PAPS (PDB 7r0u). The alignment showed a strong fit (RSMD = 0.848), indicating the accuracy of the AlphaFold2 model. Structural analysis using PyMOL revealed that the FSQQDGFY region is located on the protein’s periphery, distant from the active site, while the F56Y polymorphism is positioned near the substrate-binding pocket, 12.6 Å from vanillin and 17.9 Å from PAPS (**Figure 5d**). Although the precise functional impact of F56Y in ST4 remains to be determined, SULTs are known to metabolize plant-derived phenolic compounds (Faraji et al., 2021; Yamamoto et al., 2022), which are abundant in olive leaves (Mir-Cerdà et al., 2024).

These findings suggest that Apulian populations are genetically distinct and may be more adapted to colonizing olive trees, potentially due to local selective pressures (**Figure 5e**).

## DISCUSSION

In this study, we substantially expanded genomic resources for Cercopoidea, with a particular focus on *P. spumarius*, the primary European vector of *X. fastidiosa*. We generated new genome assemblies for *P. spumarius*, *P. tesselatus*, *P. maghresignus*, *A. alni*, and *C. vulnerata*, including a high-quality chromosome-level assembly of *P. spumarius* (12 scaffolds, 112–431 Mb). Cytogenetic and genomic analyses confirmed a 2n = 22 + XX/X0 karyotype (Kuznetsova et al., 2003) and identified scaffold 6 as the X chromosome. Supporting this, we found that chromosome 6 has lower SNP density than the autosomes, as expected for an X chromosome in an X0 system. The reduced diversity likely reflects its smaller effective population size (∼75% that of autosomes) and stronger purifying selection due to hemizygosity in males (Jaquiery et al., 2012). The absence of Y-linked loci in X0 systems may also drive the enrichment of male-biased genes on autosomes, as reported in other obligate sexual species of the Hemiptera (Li et al., 2020). Comparative analyses showed that *P. spumarius*, *P. tesselatus*, *A. alni*, and *C. vulnerata* share an X0 system, with conserved synteny across species, whereas *P. maghresignus* has an XY system and *P. italosignus* a rare XXY configuration (Maryańska-Nadachowska et al., 2012). These findings indicate that the X0 system is likely ancestral in the *Philaenus* genus, with XY systems derived, and highlight the need for additional chromosome-level assemblies to reconstruct the evolution of sex chromosome transitions in this group.

Whole-genome resequencing of 430 *P. spumarius* individuals mapped to the reference genome revealed strong population structure across Europe, with a clear isolation by distance pattern that is consistent with the species limited dispersal ability (Casarin et al., 2023). We identified five major genetic groups: (1) northern populations in Scotland, Northern Ireland, Finland, and high-altitude Pyrenees and Alps; (2) southern populations in Italy, southern France, and Portugal; (3) a distinct group in England; (4) Balkan populations extending into Turkey; and (5) a highly differentiated group in the Spanish part of the Iberian Peninsula with admixture in Portugal. Central European populations showed mixed ancestry from several clusters, except for the Iberian Peninsula, which remained distinct. Notably, individuals from southern and central Spain clustered with a specimen morphologically identified as *P. tesselatus*, showing high genetic differentiation, low heterozygosity, and evidence of long-term isolation. This pattern may reflect historical separation, possibly with origins in North Africa followed by colonization of southern Europe. Our results therefore confirm the presence of the *P. tesselatus* genotype in the Iberian Peninsula, supporting earlier observations (Maryańska-Nadachowska et al., 2010).

Despite strong nuclear divergence, the *P. tesselatus*-like populations of the Iberian Peninsula share mitochondrial haplotypes with *P. spumarius*, consistent with earlier studies reporting nearly identical mtDNA in the two species (Rodrigues et al., 2014; Seabra et al., 2020). Such mito-nuclear discordance may result from introgressive hybridization coupled with selection on mitochondrial DNA (Gompert et al., 2008). Female-biased dispersal may promote nuclear admixture while spreading maternal haplotypes (Lago et al., 2021; Casarin et al., 2023). Additionally, *Wolbachia* that is detected in the dominant northern mitochondrial haplogroup but absent in southern regions (Lis et al., 2015; Formisano et al., 2022; Kolasa et al., 2023) may drive cytoplasmic incompatibility (Shoemaker et al., 2004) favouring infected lineages and contributing to reduced nuclear heterozygosity and stronger isolation in northern and high-altitude populations. Clustering of invasive *P. spumarius* populations in the Azores, USA, and New Zealand with European populations strongly supports human-mediated dispersal. Azorean samples grouped closely with southern England, sharing mitochondrial haplotypes and nuclear components, consistent with a Northern European origin and a bottleneck-driven loss of diversity (Seabra et al., 2021). This points to southern England as the most likely source of a single colonization. In contrast, spittlebugs from New Zealand and the USA showed admixture: New Zealand and western USA populations aligned with Central Europe, while eastern USA populations clustered with Wales, suggesting multiple introductions from genetically diverse sources. All invasive populations exhibited reduced heterozygosity, consistent with founder effects (Dlugosch and Parker, 2008), underscoring the need for broader sampling to refine their origins and dispersal pathways.

We found high effective migration rates of *P. spumarius* within regions in Europe, including in southern Apulia, consistent with the rapid spread of *X. fastidiosa* in this section of Italy. Populations in Apulia, central Italy, England, and Scotland each formed distinct clusters with limited external gene flow, making our migration estimates robust. The first major outbreak of *X. fastidiosa* subsp. *pauca* strain ST53, the causal agent of OQDS, was reported near Gallipoli (province of Lecce) around 2013, with the initial spread likely beginning around 2008 and advancing northward toward Bari at an estimated rate of 10.0 km per year (95% CI: 7.5–12.5 km/year; Kottelenberg et al., 2021). This aligns with our findings of high effective migration rates for *P. spumarius* across southern Apulia. Moreover, our data show that the Apulia region is strongly connected to parts of central Italy through elevated *P. spumarius* migration, suggesting that the insect can both introduce *X. fastidiosa* into these regions and disperse within them at comparable rates. Strikingly, effective migration rates of *P. spumarius* in some regions of England and Scotland exceed those observed in Apulia, implying that if *X. fastidiosa* were introduced into these regions, its spread could outpace the 10 km/year recorded in southern Italy. Nevertheless, the UK contains geographic features such as mountain ranges and densely urbanized areas that can act as barriers to *P. spumarius* movement, thereby limiting the potential spread of the disease by these insect vectors.

We identified signatures of positive selection in genes encoding sulfotransferases (SULTs) in Apulian populations. Notably, for one SULT gene, ST4, the amino acid under selection is located within the enzyme’s catalytic domain (Toth et al., 2023), potentially influencing substrate specificity or catalytic efficiency. SULTs play diverse physiological roles in insects. In Drosophila, SULT homologs exhibit high sulphating activity toward phenolic compounds (Hattori et al., 2008). In the silkworm *Bombyx mori*, ST3 contributes to the detoxification of polyphenols, including genistein, found in mulberry leaves (Yamamoto et al., 2022). It is interesting that Spanish and Portuguese olive groves vectors were rarely found on olive trees despite their abundance in the herbaceous cover (Morente et al., 2018), whereas in the Italian orchards they were frequently associated with olives and capable of transmitting *X. fastidiosa* directly between trees (Cornara et al., 2017; Bodino et al., 2019). Moreover, olive leaves contain high levels of phenolic compounds such as oleuropein, which function as defensive metabolites against herbivores. This supports the hypothesis that *P. spumarius* populations in Apulia have adapted to olive as a host plant through selection on SULT genes, enhancing their capacity to detoxify these phenolics. Such adaptation is plausible given the dominance of olive monoculture in Apulia and the reliance on olive as the sole host during the summer, when the undergrowth has dried. SULTs have also been implicated in insecticide detoxification (Bairam et al., 2020; Xu et al., 2021), suggesting that SULT gene selection could additionally reflect adaptation to the region’s intensive insecticide use. Future functional studies will be critical to determine the specific roles of SULTs in *P. spumarius* host preference, pesticide resistance, and *X. fastidiosa* transmission.

This work provides a framework for studying vector species such as *P. spumarius* and *P. tesselatus*, and their roles in the spread of *X. fastidiosa*. If *P. spumarius* populations in Apulia have indeed evolved greater ability to colonize olive trees, this could help explain the severity of the olive outbreak in this region. In contrast, in Spain *P. spumarius* is rarely found on olive despite its abundance, and while *X. fastidiosa* is present, outbreaks there have primarily affected almond rather than olive. These findings highlight the critical importance of preventing the spread of Apulian *P. spumarius* genotypes to other olive-growing regions in Europe and North Africa. The genomic resources generated in this study provide a foundation for developing genetic diagnostic tools to identify and monitor such potentially high-risk vector populations.

## MATERIAL AND METHODS

### Genomic DNA preparation and genome sequencing

Total genomic DNA was extracted from a single adult insect of each species (see **Table S7**). The genomic DNA of the adult was extracted using Illustra Nucleon Phytopure kit according to the manufacturer’s instructions (GE Healthcare) (Wouters et al., 2020). The lysis step was performed by adding 10 µl of Proteinase K and incubating the sample in a water bath for 2 hr at 55°C. The DNA precipitation was performed using NaAc (3 m) together with isopropanol to increase the DNA yield. We assessed the quality and concentration of the DNA using nanodrop (ThermoFisher, location), and Qubit (Lifetech), and molecular weight was assessed using a FemtoPulse fragment analyser (Agilent).

PacBio library preparation and Illumina genome sequencing (HiSeq X, 150 bp paired-end) were performed by CD Genomics (New York, NY, USA) in accordance with standard protocols. 10x Chromium library preparation was performed by Novogene Bioinformatics Technology Co (Beijing, China) in accordance with standard protocols. For *P. spumarius* and *A. alni* a pool of mixed-stage samples was used to construct a Hi-C chromatin contact map to enable a chromosome-level assembly. Phase Genomics, Seattle, US created the Hi-C library with the DpnII restriction enzyme following a similar protocol to Lieberman-Aiden et al. (2009). The Hi-C library was sequenced on an Illumina (San Diego, CA, USA) HiSeq X sequencer and 150-bp paired-end reads were generated.

### Genome assembly of *Philaenus spumarius*

PacBio reads were first assembled using two de novo assemblers: flye v2.7.1 (Kolmogorov et al., 2019) and wtdbg2 (Ruan and Li, 2019). Both assemblies were firstly polished by pilon v1.23 (Walker et al., 2014) using Illumina reads and then merged using quickmerge v0.3 (Chakraborty et al., 2016), aiming to maximise genome completeness and minimise duplicated regions caused by under-collapsed heterozygosity. Redundant haplotigs were removed in purge_dups (Guan et al., 2020). The assembly was improved by performing iterative scaffolding using 10X Genomics raw data (from Biello et al., 2020) following the procedure set out in Biello et al. (2021). Briefly, we performed two rounds of Scaff10x (https://github.com/wtsi-hpag/Scaff10X) with the parameters “-longread 1 -edge 50000 - block 50000,” followed by misassembly detection and correction with tigmint v1.1.2 (Jackman et al., 2018). These steps were followed by a final round of scaffolding with Assembly Round-up by Chromium Scaffolding (arcs) (Yeo et al., 2018).

We aligned the Hi-C reads to the assembly using the juicer pipeline (Durand et al., 2016). The assembly was then scaffolded with Hi-C data into chromosome-level organization using the 3d-dna pipeline (Dudchenko et al., 2017), followed by manual correction using Juicebox Assembly Tools (jbat) (Dudchenko et al., 2018). The assembly was polished after jbat review using the 3d-dna seal module to reintegrate genomic content removed from super-scaffolds through false-positive manual editing to create a final scaffolded assembly.

### Genome assembly of Philaenus tesselatus and P. maghresignus

PacBio reads were first assembled using two de novo assemblers: Flye v2.9 (Kolmogorov et al., 2019) and wtdbg2 (Ruan and Li 2019). Both assemblies were firstly polished by Polca (Zimin and Salzberg, 2020) using Illumina reads and then merged using quickmerge v0.3 (Chakraborty et al., 2016), aiming to maximise genome completeness and minimise duplicated regions caused by under-collapsed heterozygosity. Redundant haplotigs were removed in purge_dups (Guan et al., 2020).

### Genome assembly of *Aphrophora alni*

PacBio reads were first assembled using two de novo assemblers: Flye v2.9 (Kolmogorov et al., 2019) and wtdbg2 (Ruan and Li, 2019). Both assemblies were firstly polished by Polca (Zimin and Salzberg, 2020) using Illumina reads and then merged using quickmerge v0.3 (Chakraborty et al., 2016), aiming to maximise genome completeness and minimise duplicated regions caused by under-collapsed heterozygosity. Redundant haplotigs were removed in purge_dups (Guan et al., 2020).

We aligned the Hi-C reads to the assembly using the juicer pipeline (Durand et al., 2016). The assembly was then scaffolded with Hi-C data into chromosome-level organization using the 3d-dna pipeline (Dudchenko et al., 2017), followed by manual correction using Juicebox Assembly Tools (jbat) (Dudchenko et al., 2018). The assembly was polished after jbat review using the 3d-dna seal module to reintegrate genomic content removed from super-scaffolds through false-positive manual editing to create a final scaffolded assembly.

### Genome assembly of *Cercopis vulnerata*

To create the de novo assembly, the 10X Genomics linked-read data were assembled using supernova 2.1.1 (Weisenfeld et al., 2017) with the default parameters and the recommended number of reads (--maxreads = 199222871) to produce the pseudohaplotype assembly output (--style = pseudohap). We improved the initial supernova assembly by performing iterative scaffolding using all of the 10X Genomics raw data following the procedure set out in Biello et al. (2021). Briefly, we performed two rounds of Scaff10x (https://github.com/wtsi-hpag/Scaff10X) with the parameters “-longread 1 -edge 50000 - block 50000,” followed by misassembly detection and correction with tigmint (Jackman et al., 2018).

### Quality assessment

We checked each assembly for contamination using the blobtools pipeline v1.0.1 (Kumar et al., 2013; Laetsch and Blaxter, 2017) by generating taxon annotated GC content-coverage plots (known as “BlobPlots”). Each scaffold was annotated with taxonomy information based on blastn (Basic Local Alignment Search Tool) v2.2.31 (Camacho et al., 2009) searches against the National Center for Biotechnology Information (NCBI) nucleotide database (nt, downloaded April 2, 2019) with the options “-outfmt ’6 qseqid staxids bitscore std sscinames sskingdoms stitle’ - culling_limit 5 -evalue 1e-25.” To calculate average coverage per scaffold, we mapped the Illumina raw reads to the assembly using bwa-mem v0.7.17 (Burrow-Wheeler Aligner) (Li, 2013) with default parameters. The resulting BAM file was sorted with samtools v1.9 (Li et al., 2009) and passed to blobtools along with the table of blastn results.

A frozen release was generated for the final assembly of each species with scaffolds renamed and ordered by size with seqkit v0.12.0 (Shen et al., 2016). We assessed the quality of the genome assembly by searching for conserved, single copy, Hemiptera genes (n = 2,510) with Benchmarking Universal Single-Copy Orthologs (BUSCO) v5.1.2 (Manni et al., 2021) and by analysis of k-mer spectra with Merqury v1.3 (Rhie et al., 2020) to compare k-mer content of the raw sequencing reads to the k-mer content of the assembly.

### Mitochondrial assembly

Mitochondrial genomes of *P. spumarius*, *P. tesselatus*, *P. maghresignus*, *A. alni* and *C. vulnerata* were assembled from short reads using MitoZ v2.3 (Meng et al., 2019) with options “--genetic_code 5 --clade Arthropoda --fastq_read_length 150 --insert_size 250 --run_mode 2 --filter_taxa_method 1 --requiring_taxa ’Arthropoda’” for Arthropoda species. All mitochondrial genomes were annotated using MitoZ with the function annotate.

### Repeat annotation

Two methods were combined to identify the repeat contents in the genome of *P. spumarius*, *P. tesselatus*, *P. maghresignus*, *A. alni*, *C. vulnerata* and *C. versicolor*: homology-based and de novo prediction. Using the homology-based analysis, we identified the known transposable elements (TE) within the genome using RepeatMasker v4.0.7 (Tarailo-Graovac and Chen, 2009) using the RepBase Insecta repeat library (Bao et al., 2015) with the parameters “-e ncbi -species insecta -a -xsmall -gff” (Jurka et al., 2005). For de novo prediction, we constructed a de novo repeat library of each genome using RepeatModeler v2.0.2 (Flynn et al., 2020). Finally, we merged the library files of the two methods and used RepeatMasker to identify the repeat contents.

### Genome annotation

Total RNA was extracted from three pools of *P. spumarius*. Two containing adults (females and males, respectively) and one containing nymphs, collected from the same location (**Table S7**). For *P. tesselatus*, *P. maghresignus*, *A. alni* and *C. vulnerata,* total RNA was extracted from two pools containing adults (females and males, respectively). For *C. versicolor*, RNAseq data was downloaded by NCBI (Chen et al., 2022). The sample was ground under liquid nitrogen in a 1.5-ml Eppendorf tube using a plastic pestle. RNA was extracted using Trizol (Sigma) according to the manufacturer’s protocol. RNA was further purified using RNeasy with on-column DNAse treatment (Qiagen) according to the manufacturer’s protocol and eluted in 100 ml of nuclease-free water. RNA quality was assessed by electrophoresis of 5 μl denatured in formamide on a 1% agarose gel. Purity was assessed using a Nanodrop spectrophotometer (ThermoFisher) to measure the A260/A280 and A260/A230 ratios. Concentration of RNA was measured, and the presence of contaminating DNA was assessed using a Qubit (Lifetech). Library prep and stranded RNAseq was done by Novogene, yeilding 75 millions of reads per library.

For the gene prediction, we first mapped the quality- and adapter-trimmed RNA-seq reads to the soft-masked assembly with hisat2 v2.1.0 (Kim et al., 2015), followed by sorting with samtools v1.10 (Li et al., 2009). Quality control and trimming for adapters and low-quality bases (quality score <20) of the RNA-seq raw reads were performed using fastqc v0.11.8 (Andrews, 2010) and TrimGalore v0.5.0 (https://github.com/FelixKrueger/TrimGalore), respectively. All the BAM files were filtered to remove invalid splice junctions with Portcullis v1.1.2 (Mapleson et al., 2018). For the prediction of gene loci and structures, we used the BRAKER3 pipeline (Gabriel et al., 2024). It uses the softmasked genome, filtered RNA-seq alignments and OrthoDB10 protein data (Kuznetsov et al., 2023) (https://bioinf.uni-greifswald.de/bioinf/partitioned_odb11/) as input to subsequently run the gene prediction tool GeneMark-ETP v1.00 (Brůna et al., 2024) and then AUGUSTUS v3.4.0 (Stanke et al., 2008) for its annotation process. The BRAKER transcript selector, TSEBRA v1.0.3 (Gabriel et al., 2021), was employed to unify predictions from these 3 sources and configured such that RNA evidence had greater weight than protein evidence. To assess the completeness of the annotation, we used BUSCO v5.1.2 (Manni et al., 2021) using the Hemiptera odb10 reference database in protein mode. Protein domains were annotated by searching the InterPro v32.0 (Hunter et al., 2012) and Pfam v27.0 (Punta et al., 2012) databases, using Interproscan v5.52 (Quevillon et al., 2005) and hmmer v3.3 (Finn et al., 2011), respectively.

### Genetic Sexing from Coverage

The resequencing reads from three individuals of each sex were mapped to the reference genome using bwa-mem v0.7.17 (Burrow-Wheeler Aligner) (Li, 2013) with default parameters. Samtools v1.9 (Li et al., 2009) was used to sort and index the mapped reads, and coverage depth across the genome was calculated using the samtools depth command. The coverage depth data for the male and female individuals were compared across all scaffolds. For each scaffold, the mean coverage was calculated separately for the male and female individuals. The ratios of male-to-female coverage were computed for each scaffold. Coverage depth differences were visualized using R (R Core Team citation) to plot male and female coverage across the genome.

### Phylogeny and comparative genomics

Orthologous groups in Auchenorrhyncha, genomes were identified from the predicted protein sequences of *P. spumarius*, *P. tesselatus*, *P. maghresignus*, *A. alni*, *C. vulnerata* and ten other Auchenorrhyncha genomes already published (see **Table S3**). As an outgroup, we included the genomes of two aphids: *Myzus persicae* (Mathers et al., 2021; Liu et al., 2024) and *Eriosoma lanigerum* (Biello et al., 2021) (**Table S3**). We used the longest transcript to represent the gene model when several transcripts of a gene were annotated. Orthofinder v2.5.4 (Emms and Kelly, 2019) with diamond v0.9.14 (Buchfink et al., 2015), Multiple Alignment using Fast Fourier Transform (MAFFT) v7.305 (Katoh and Standley, 2013) and with RAxML v8.2.12 (Stamatakis, 2014) were used to cluster proteins into orthogroups, reconstruct gene trees and estimate the species tree

### Synteny analysis

Syntenic blocks of genes in Cercopoidea were identified between the chromosome-level genome assemblies of *P. spumarius*, *Aphrophora alni* and *Callitettix versicolor* (Chen et al., 2022) (see **Table S3**) using Genespace v1.2.3 (Lovell et al., 2022), which runs OrthoFinder and McScanX to identify and visualize syntenic relationships (should also cite R here probably). As outgroup we used the genome of *Homalodisca vitripennis* (GCF_021130785.1). Riparian plot was obtained using the synteny calculation blocks obtained from Genespace.

### Re-sequencing sample preparation, mapping, and variant calling

We collected a geographically dispersed set of 2,839 *P. spumarius* samples, selected to maximize information about genomic differentiation across the species range (**Table S8, Figure S7**).

Samples were collected in 2019–2021 from locations that cover broad regions throughout Europe, and samples from non-native regions such as the US, New Zealand and the Azores (**Table S8, Figure S7**). Within each location, nymphs or adults were collected from different plant hosts. Field-collected samples were morphologically inspected by taxonomists. Libraries were sequenced on a HiSeq X using a 150 bp paired-end read metric to an average coverage of ∼12x. FastQC was used to check the quality of the raw reads obtained and reads were trimmed using TrimGalore (https://github.com/FelixKrueger/TrimGalore). To call variants, data were first aligned to the chromosome-scale assembly of Phspu_JIC_v2 assembly using BWA-MEM2 (Vasimuddin et al., 2019). SAM format files were sorted, indexed and converted into BAM files using SAMtools v1.10 (Danecek et al., 2021). GATK (McKenna et al., 2010) and picardtools (http://broadinstitute.github.io/picard) were used to identify and realign reads around indels (RealignerTargetCreator, IndelRealigner) as well as remove duplicates (MarkDuplicates, all default settings). BCFtools v1.10.2 (Danecek et al., 2021) mpileup and call function were used to identify SNPs in each sample. VCFtools v.0.1.16 (Danecek et al., 2011) was used to filter the dataset to only retain biallelic SNPs with SNP quality scores (QUAL) ≥30 with the option “--remove-indels --max-alleles 2 --max-missing 0.9 --minQ 30 --min-meanDP 5 --max-meanDP 35 --minDP 5 --maxDP 35”. Finally, all sites included in repeated genomic regions or in regions with a low mappability score, calculated using the GEM algorithm (Derrien et al., 2012) and setting a maximum mismatch of 0.04 in a 150 bp read, were removed.

Raw sequencing reads in FASTQ format were mapped to the mitochondrial reference genome using BWA-MEM2 (Vasimuddin et al., 2019) with default parameters. The resulting SAM file was converted to BAM format and sorted using SAMtools v1.10 (Danecek et al., 2021). To reduce potential biases from PCR amplification, duplicates were marked using Picard Tools MarkDuplicates (http://broadinstitute.github.io/picard). Variants were identified using GATK v4.1.4.0 (McKenna et al., 2010) HaplotypeCaller in GVCF mode, with the ploidy set to 1 to account for the haploid nature of mitochondrial DNA. The resulting GVCF file was filtered to retain only high-quality variants based on standard quality metrics. The filtered VCF file was then processed using BCFtools v1.10.2 (Danecek et al., 2021) consensus to generate the final consensus sequence in FASTA format.

### Population structure

Whole genome mitochondrial sequences were aligned using MUSCLE v3.8 (Edgar, 2004), and a phylogenetic tree was constructed using maximum likelihood estimation with IQ-TREE v1.6.10 (Nguyen et al., 2015). The tree was edited in FigTree v1.4.4 (http://tree.bio.ed.ac.uk/software/figtree/) for improved visualization of sample relationships and their overall clustering patterns. P. maghresignus was used as the outgroup for the phylogenetic analysis.

To further investigate the population structure using whole-genome sequencing data, we applied several complementary methods. Principal Component Analysis (PCA) was performed using PLINK2 (Purcell et al., 2007), after pruning for linkage disequilibrium (LD) with the "indep-pairwise" function, using the parameters (–indep-pairwise 50 5 0.1).

We also conducted ADMIXTURE analysis to estimate the maximum likelihood of individual genetic ancestries from multiple SNP datasets. For this analysis, the same set of SNPs pruned using PLINK v2 (Chang et al., 2015) as in the PCA clustering was used. ADMIXTURE v1.3 (Alexander et al., 2009) was used to analyze the population structure and infer potential ancestral populations. The most probable number of ancestral populations (K) was selected based on the lowest cross-validation error after testing a range of K values (1–8) with 10 independent replicates and a bootstrap of 100.

To explore the genetic relationships further, a phylogenetic network was constructed using SPLITSTREE v4.14.6 (Huson and Bryant, 2006) and the Neighbor-net algorithm (Bryant and Moulton, 2004).

Finally, FST, inbreeding statistics (FIS), and heterozygosity were calculated using VCFtools v.0.1.16 (Danecek et al., 2011) across non-overlapping 50 kb windows.

### Isolation-by-Distance and effective migration surfaces

We identified corridors and barriers to gene flow in the native range of *P. spumarius* using the fEEMS v1.0.1 (Marcus et al., 2021) software, that estimates effective migration surfaces across space. We adopted a grid size of 100 km^2^ that was the best compromise between computational burden and model misspecification. All populations were assigned to 54 different grids. Individuals from different sampling locations within the same grid that were genetically close were grouped, leaving a minor risk of bias in migration surface estimation with adjacent grids.

Connectivity on a local scale in Italy and the UK was explored by running EEMS v1 (Petkova et al., 2016) which assesses the decay of genetic similarity between sampling locations under a null model of isolation by distance. We created an average genetic dissimilarity matrix using the bed2diffs function as is implemented in EEMS (Petkova et al., 2016). EEMS was run over 2,000 demes, for 2 million generations, discarding the first 1 million as burnin. The results were visualized using rEEMSplots.

We also tested for isolation by distance or IBD (Wright, 1943) by conducting a linear regression of linearized pairwise F_ST_, or F_ST_/(1−F_ST_), against the Euclidean distance between populations in Italy (including the main islands) and the UK. Pairwise F_ST_ values were calculated using VCFtools v.0.1.16 (Danecek et al., 2011), across non-overlapping 50 kb windows with a sliding window step size of 25 kb. We estimated the geographic distances among populations as Euclidean distances, using the “dist” function from the R package stats v3.3.1 (R Core Team, 2013). The statistical significance of IBD was evaluated with a Mantel test (Pearson’s product moment correlation; 9,999 permutations) using the R package vegan v2.6.4 (Oksanen et al., 2020).

### Selection scans

To detect selection in the Apulia population, we used Population Branch Statistics (PBS) with data from four populations, as described by Jiang et al. (2020). We focused on genomic regions specifically differentiated in Apulia compared to Northern Italy, Norfolk (UK), and Spain. These three populations were chosen due to having the highest number of samples (>20 samples per population). Mean pairwise FST values were calculated within non-overlapping 50 kb windows using VCFtools v.0.1.16 (Danecek et al., 2011). Negative FST values were set to zero, and fixed differences were set to 0.99 to avoid infinite branch lengths. Finally, we applied the Cavalli-Sforza transformation and calculated the PBS values using the following equation:

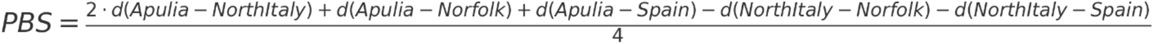

Where d is the log-transformed distance between two populations. The windows showing a negative PBS, which can be obtained in case of small sample size, were set to zero as they do not have a biological meaning.

### RT-PCR and sequencing

RNA was extracted from *P. spumarius* collected from UK, Portugal, and Apulia (Italy), and first strand cDNA was synthesised using Superscript IV (Life Technologies) primed with the St4-CDS-R primer (TCATACTAAGGGAAATCTCAAATCG). PCR amplification using Phusion (NEB) with the St4-CDS-F (ATGGATCACAATATAAATGAGG) and St4-CDS-R amplified a product of approximately 1kb. Purified PCR products were sequenced (Azenture Genewiz) using the forward and reverse primers, confirming the ST4 CDS as 948bp spanning the two gene models.

### Structural modelling

The ST4 protein structure was modelled using Alphafold2 (Jumper et al., 2021), with default settings, and the highest PLDDT scoring model (see **Figure S15** for PLDDT plots) was searched against the PDB database using Foldseek (van Kempen et al., 2024), which identified the *Anopholes gambiae* sulphotransferase (AgST: 7r0o) as highly similar (Seq ID 41.1, e<1E-33). Structures were visualised and aligned using PyMol (Yuan et al., 2017).

### Data Availability

The genome raw reads and the genome assemblies are available at the National Center for Biotechnology Information (NCBI) with the BioProject no. PRJNA1299764 and PRJNA1304604. Analysis scripts are publicly available on GitHub (https://github.com/rsbiello/Spittlebugs_popgen).

## Acknowledgements

We thank Matteo Gravino, Yazhou Chen, Adi Kliot, Rebecca Corkill, Sylvain Capdevielle and Weijie Huang for help with DNA extractions from insect samples. We are also grateful to Hernán A. Burbano and Sergio M. Latorre (University College London) for their feedback on various drafts of the manuscript, and the JIC Molecular Genetics, Research Computing, and Entomology platforms for their support. The authors used ChatGPT to assist with grammar and clarity during manuscript editing. All text was reviewed, revised, and verified by the authors, who take full responsibility for the content.

Members of the Spittlebug Collection Consortium (with ORCID where applicable) are Emma J. Back, John S. Badmin, Lisa Blackburn, Domenico Bosco (0000-0003-3945-2752), Rebecca Cairns, Brian A. Carter (0000-0003-1342-0197), Simon Charlesworth, Debbie A. Collins, Larissa E. Collins (0000-0001-5449-0015), Simon Conyers, Gerard Clover, Damian J. De Marzo, Rachel E. Down, Anne Edwards (0000-0002-3720-0968), Alberto Fereres (0000-0001-6012-3270), Christopher Foster (0000-0002-7635-6797), Susannah Gill, Massimo Giorgini (0000-0001-8670-0945), Helena Glassup, Caroline Gorton, Karl Graham, Claire Harkin (0000-0002-4106-038X), Alvin J. Helden (0000-0002-8607-8356), Fiona Highet (0000-0002-0611-7723), Tim Howell, Stephen J. Jenkins (0000-0002-0233-5424), Abigail Jenkins, Daniel Jenkins, Eleanor P. Jones (0000-0001-7833-7226), Anna Jordan, Jelena Jović (0000-0002-7623-0553), Aliesha Kean (0000-0003-2284-5592), Kirsten Knox (0000-0002-5487-1763), Katherine Lester (0000-0002-1534-8590), Otto Lindenbaum, Sam Lindenbaum, Kirsty Lindenbaum, Rodney Monteith, Gabriele Moro (0009-0007-9971-006X), Sandra Åhlén Mulio (0000-0002-0272-1229), Miranda Mugford, Kieran Murphy, Ana Pérez-Sierra (0000-0001-5403-1433), Chris R. J. Pollard (0000-0003-1278-6891), Jessica Prickett, Maria Teresa Rebelo (0000-0002-2724-2195), Verena Rösch (0000-0002-0662-4338), Sofia G. Seabra (0000-0003-1413-2349), Martina Šeruga-Musić (0000-0002-0524-0834), Victor Soria-Carrasco (0000-0002-6568-5544), Alan J. A. Stewart (0000-0001-7878-8879), Jean-Claude Streito (0000-0002-9104-2657), Jason Sumner-Kalkun (0000-0002-6485-1731), D. John I. Thomas, Vinton Thompson (0000-0003-3257-0107), Sietse van der Linde (0000-0002-1255-8963), Jessica Vereijssen (0000-0002-2095-6312), Joana G. Vicente (0000-0001-8442-5935), Diana Westmoreland, Michael R. Wilson, Roland H. M. Wouters (0000-0002-2380-4875), and Emma Louise Wright.

## Funding

This project was funded by the BRIGIT project (BB/S016325/1, to SAH) through a grant from the UK Research and Innovation (UKRI) under the Strategic Priorities Fund, in collaboration with the Biotechnology and Biological Sciences Research Council (BBSRC) with support from the Department for Environment, Food and Rural Affairs (DEFRA) and the Scottish Government. Additional support was provided by BBSRC via the Institute Strategy Programmes (BBS/E/J/000PR9797, BBS/E/J/000PR9798, BBS/E/JI/230001B and BBS/E/JI/230001D, to SAH) awarded to the John Innes Centre (JIC), which is grant-aided by the John Innes Foundation.

### Author Contributions

**Conceptualization**: Roberto Biello, Saskia A. Hogenhout, The Spittlebug Consortium Alan J. A. Stewart.

**Formal analysis**: Roberto Biello, Sam T. Mugford, Saskia A. Hogenhout.

**Funding acquisition**: Saskia A. Hogenhout.

**Investigation**: Roberto Biello, Sam T. Mugford, Qun Liu, Saskia A. Hogenhout, The Spittlebug Consortium All.

**Methodology**: Roberto Biello, Sam T. Mugford, Thomas C. Mathers, Qun Liu, The Spittlebug Consortium Gerard Clover, Fiona Highet, Alan J. A. Stewart, Michael Wilson.

**Project administration**: Saskia A. Hogenhout, The Spittlebug Consortium Gerard Clover, Larissa E. Collins, Fiona Highet, Alan J. A. Stewart, Michael Wilson.

**Resources**: Roberto Biello, Sam T. Mugford, Qun Liu, Saskia A. Hogenhout, The Spittlebug Consortium All.

**Software**: Roberto Biello, Thomas C. Mathers.

**Supervision**: Sam T. Mugford, Thomas C. Mathers, Qun Liu, Saskia A. Hogenhout, The Spittlebug Consortium Larissa E. Collins.

**Validation**: Roberto Biello, Sam T. Mugford, Thomas C. Mathers, Qun Liu, Saskia A. Hogenhout.

**Visualization**: Roberto Biello, Sam T. Mugford, Saskia A. Hogenhout, The Spittlebug Consortium Victor Soria-Carrasco.

**Writing - original draft**: Roberto Biello, Sam T. Mugford, Saskia A. Hogenhout.

**Writing - review & editing**: Roberto Biello, Sam T. Mugford, Thomas C. Mathers, Qun Liu, Saskia A. Hogenhout, The Spittlebug Consortium Domenico Bosco, Tim Howell, Eleanor P. Jones, Sofia G. Seabra.

Competing Interest

The authors declare that they have no competing interests.

---

**Figure S1.**
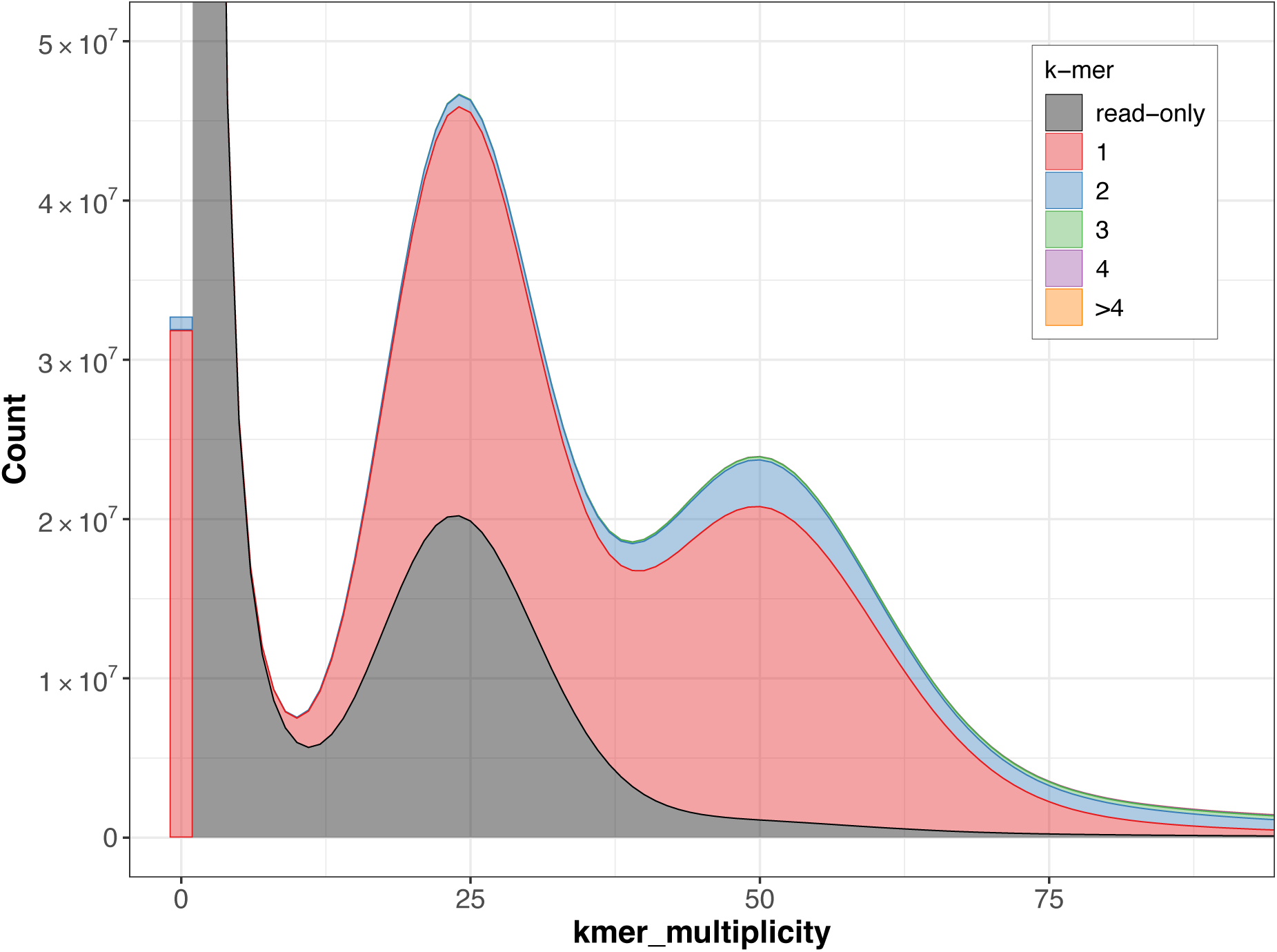
Merqury k-mer plots assessing k-mer content of Illumina raw reads used for generating the genome assembly of *Philaenus spumarius* (Phspu_JIC_v2) described herein. The black area of the graphs represents the distribution of k-mers present in the reads but not in the assembly and the red area represents the distribution of k-mers present in the reads and once in the assembly. Other colours show k-mers found multiple times in the genome assembly.

**Figure S2.**
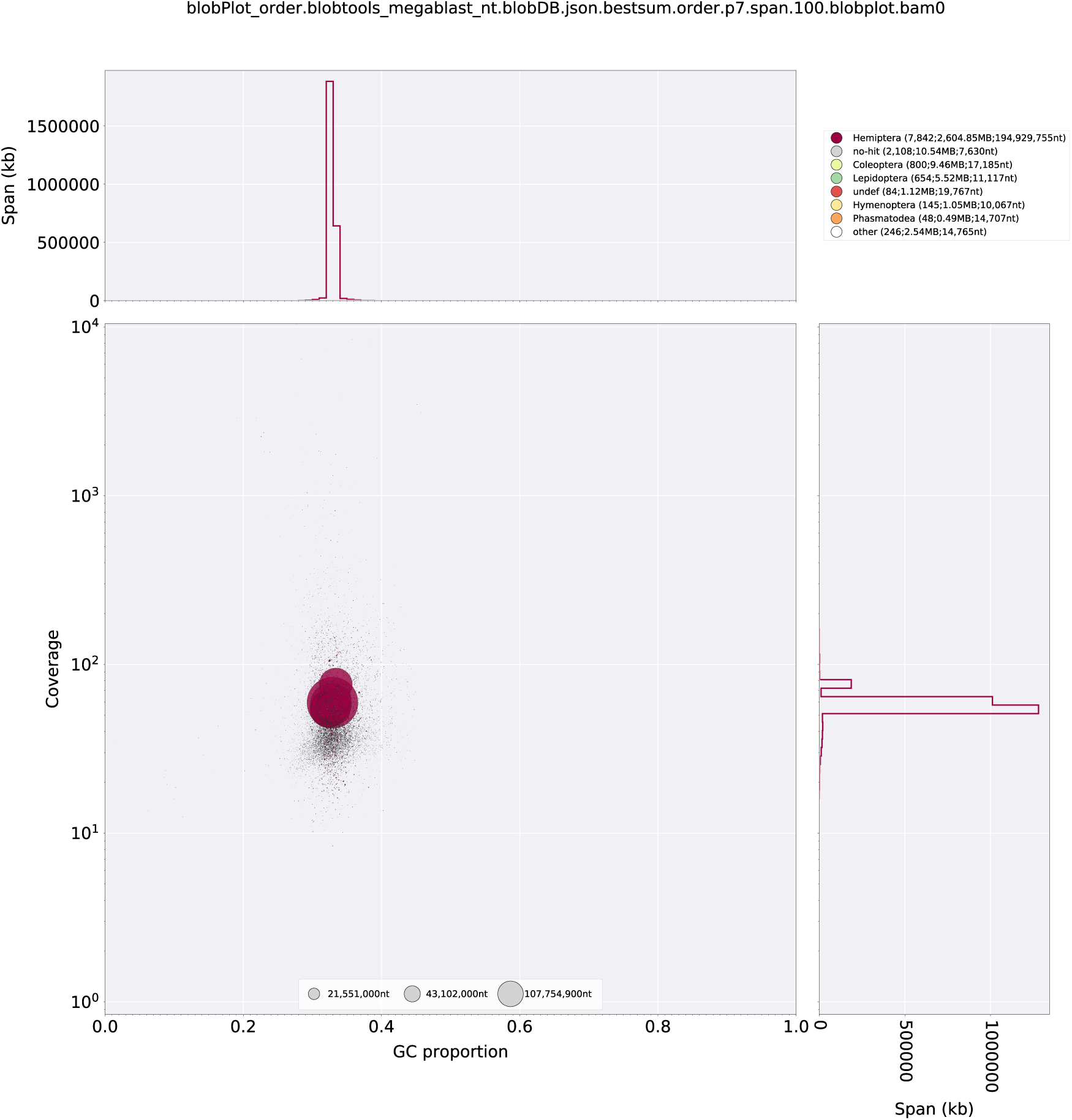
Taxon-annotated GC content-coverage plot of Phspu_JIC_v2 genome assembly scaffolds. Each circle represents a scaffold in the assembly, scaled by length, and coloured by order-level NCBI taxonomy assigned by BlobTools. The X axis corresponds to the average GC content of each scaffold and the Y axis corresponds to the average coverage based on alignment of raw Illumina reads. Marginal histograms show cumulative genome content (in Kb) for bins of coverage (Y axis) and GC content (X axis).

**Figure S3.**
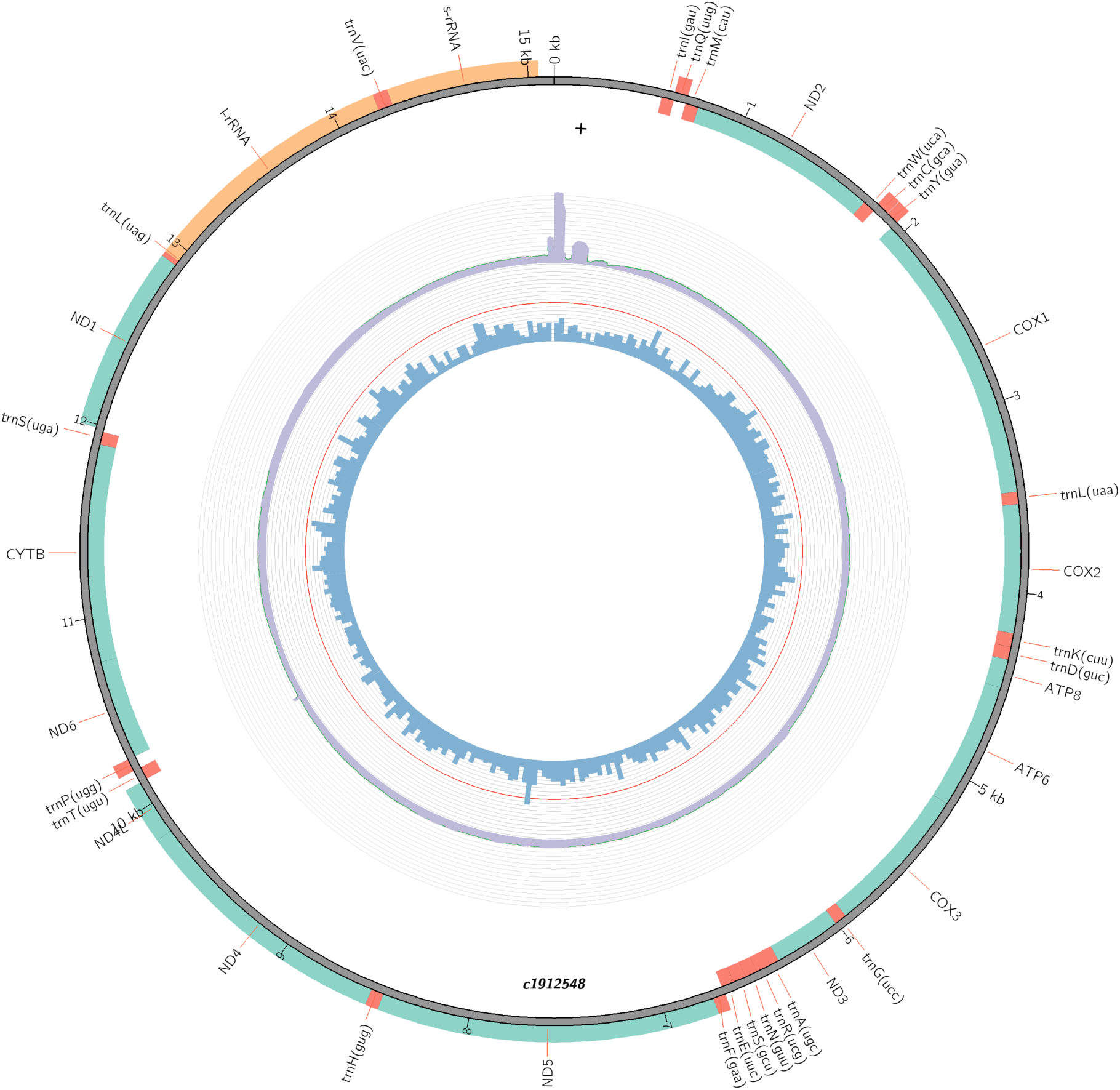
Mitochondrial genome of *Philaenus spumarius*. The circular mitochondrial genome comprises 38 genes typical of insect mitochondria, including 13 protein-coding genes, two ribosomal RNA genes (rRNAs), 23 transfer RNA genes (tRNAs), and an A+T-rich control region.

**Figure S4.**
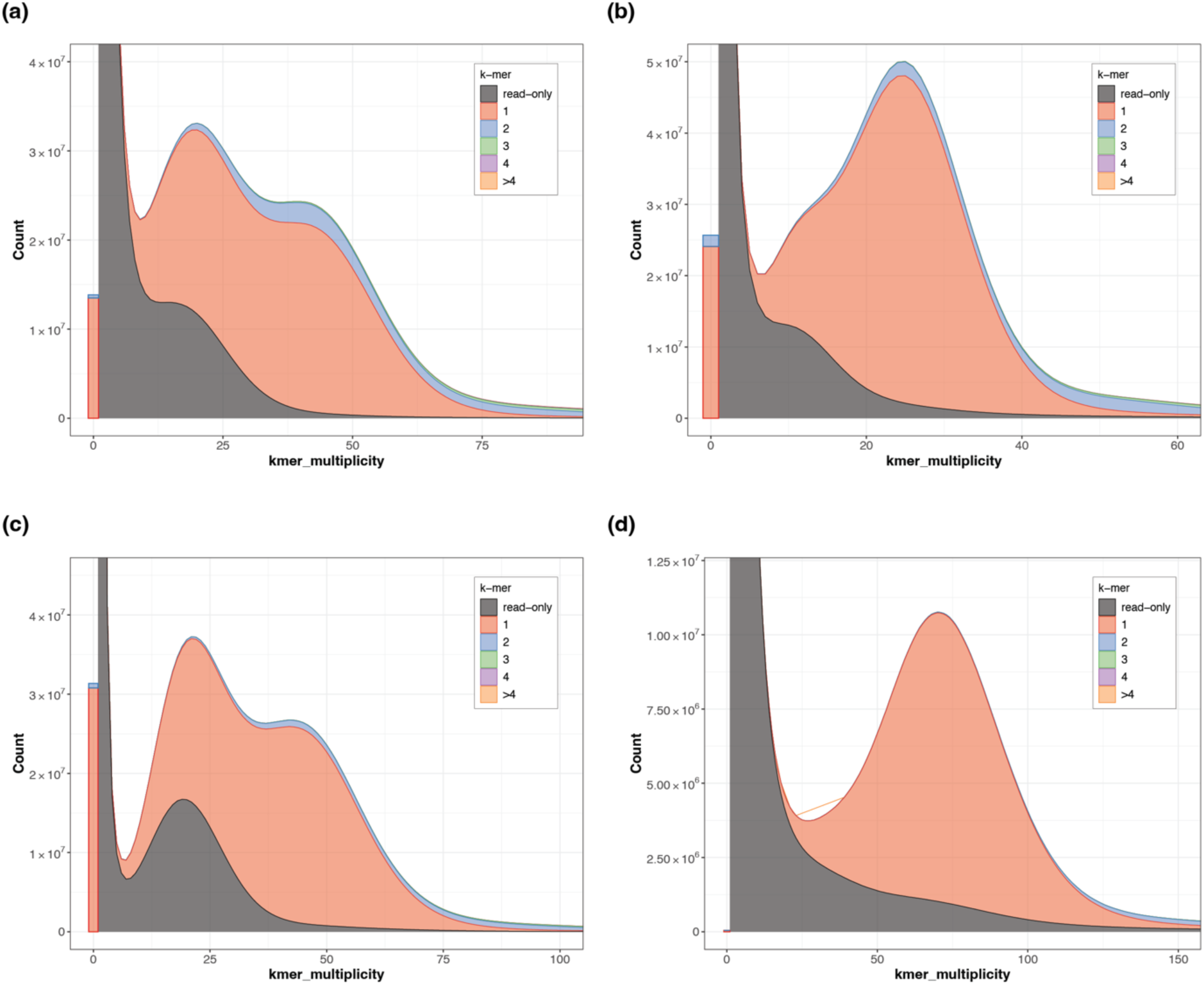
Merqury k-mer plots assessing k-mer content of Illumina raw reads used for generating the genome assembly of (a) *Philaenus tesselatus,* (b) *P. maghresignus*, **(c) *Aphrophora alni* and (d) *Cercopis vulnerata*.** The black area of the graphs represents the distribution of k-mers present in the reads but not in the assembly and the red area represents the distribution of k-mers present in the reads and once in the assembly. Other colours show k-mers found multiple times in the genome assembly.

**Figure S5.**
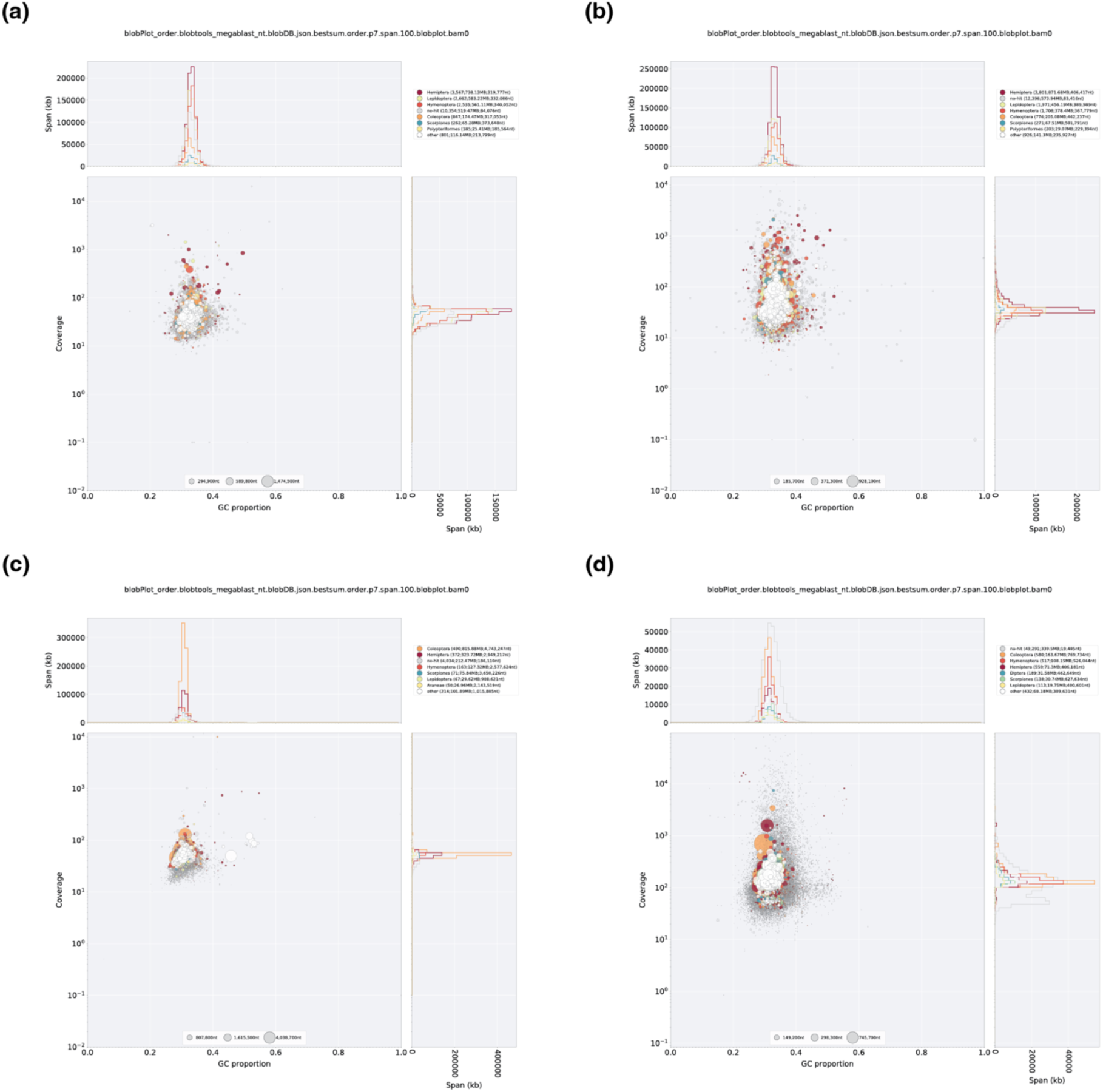
Taxon-annotated GC content-coverage plot of genome assembly scaffolds of (a) *Philaenus tesselatus,* (b) *P. maghresignus*, (c) *Aphrophora alni* and (d) *Cercopis vulnerata*. Each circle represents a scaffold in the assembly, scaled by length, and coloured by order-level NCBI taxonomy assigned by BlobTools. The X axis corresponds to the average GC content of each scaffold and the Y axis corresponds to the average coverage based on alignment of raw Illumina reads. Marginal histograms show cumulative genome content (in Kb) for bins of coverage (Y axis) and GC content (X axis).

**Figure S6.**
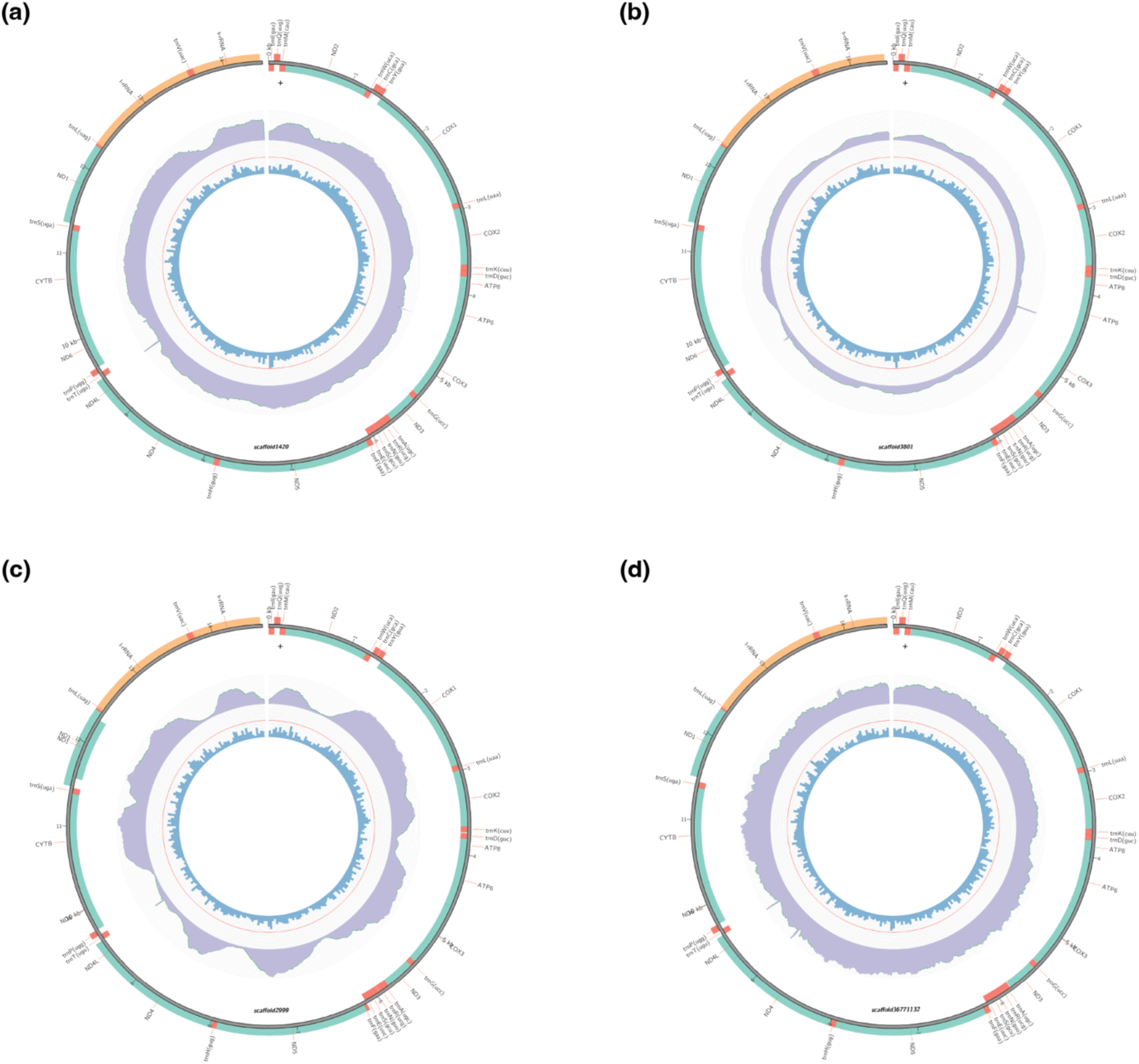
Mitochondrial genomes of (a) *Philaenus tesselatus,* (b) *P. maghresignus*, (c) *Aphrophora alni* and (d) *Cercopis vulnerata*. The circular mitochondrial genome comprises 38 genes typical of insect mitochondria, including 13 protein-coding genes, two ribosomal RNA genes (rRNAs), 23 transfer RNA genes (tRNAs), and an A+T-rich control region.

**Figure S7.**
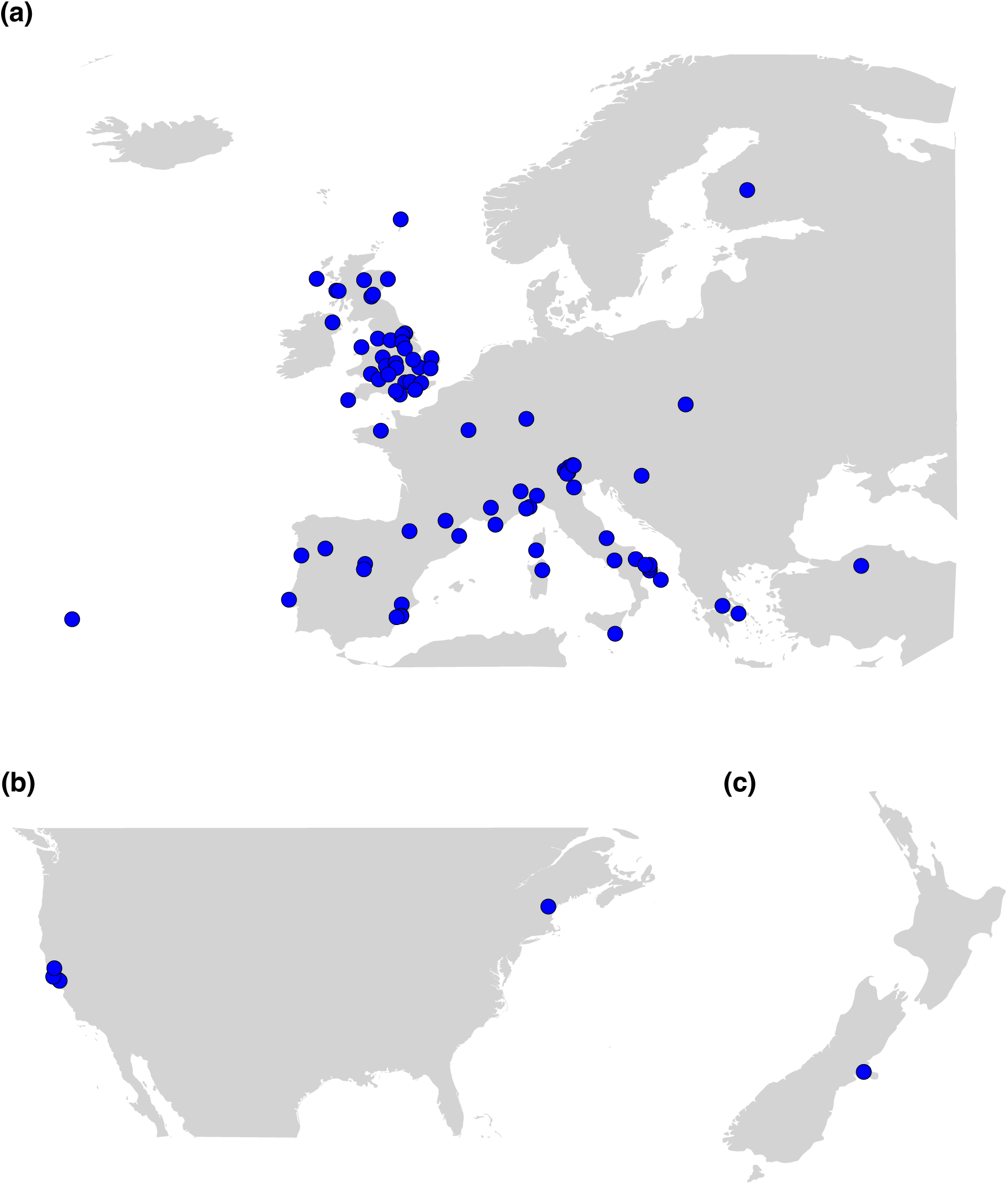
Collection sites of individuals used for *Philaenus spumarius* populaton stucture analyses. At each location ∼4 individuals were processed for DNA extraction and genome resequencing (**a**) Europe, (**b**) North America, (**c**) New Zealand.

**Figure S8.**
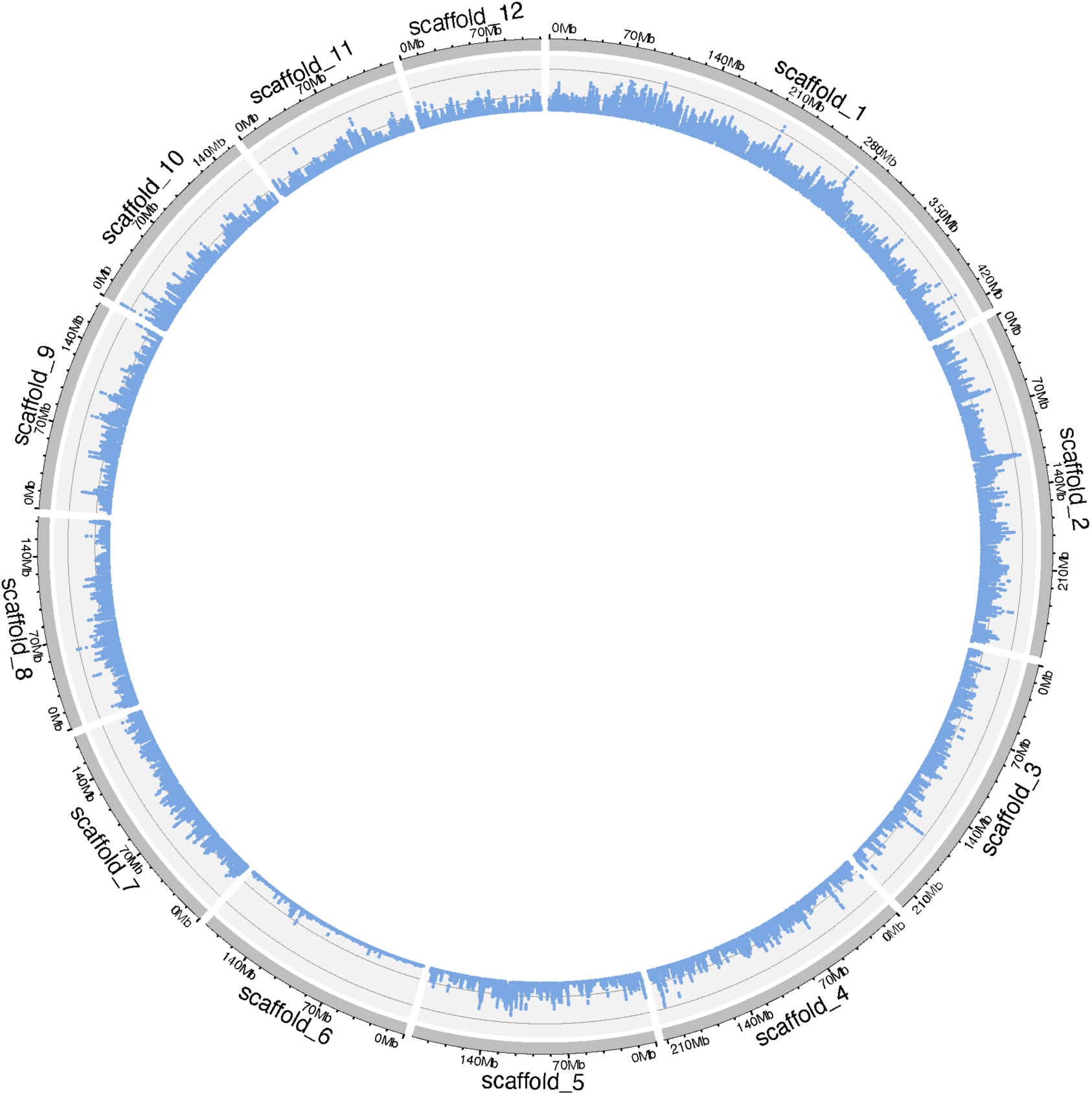
Circos plot showing the chromosomal distribution of single nucleotide polymorphisms (SNPs) identified from whole-genome resequencing data of 430 *Philaenus spumarius* individuals, mapped to the chromosome-level *P. spumarius* genome assembly (Phspu_JIC_v2). Each dot represents the SNP density calculated within a 50-kb genomic window.

**Figure S9.**
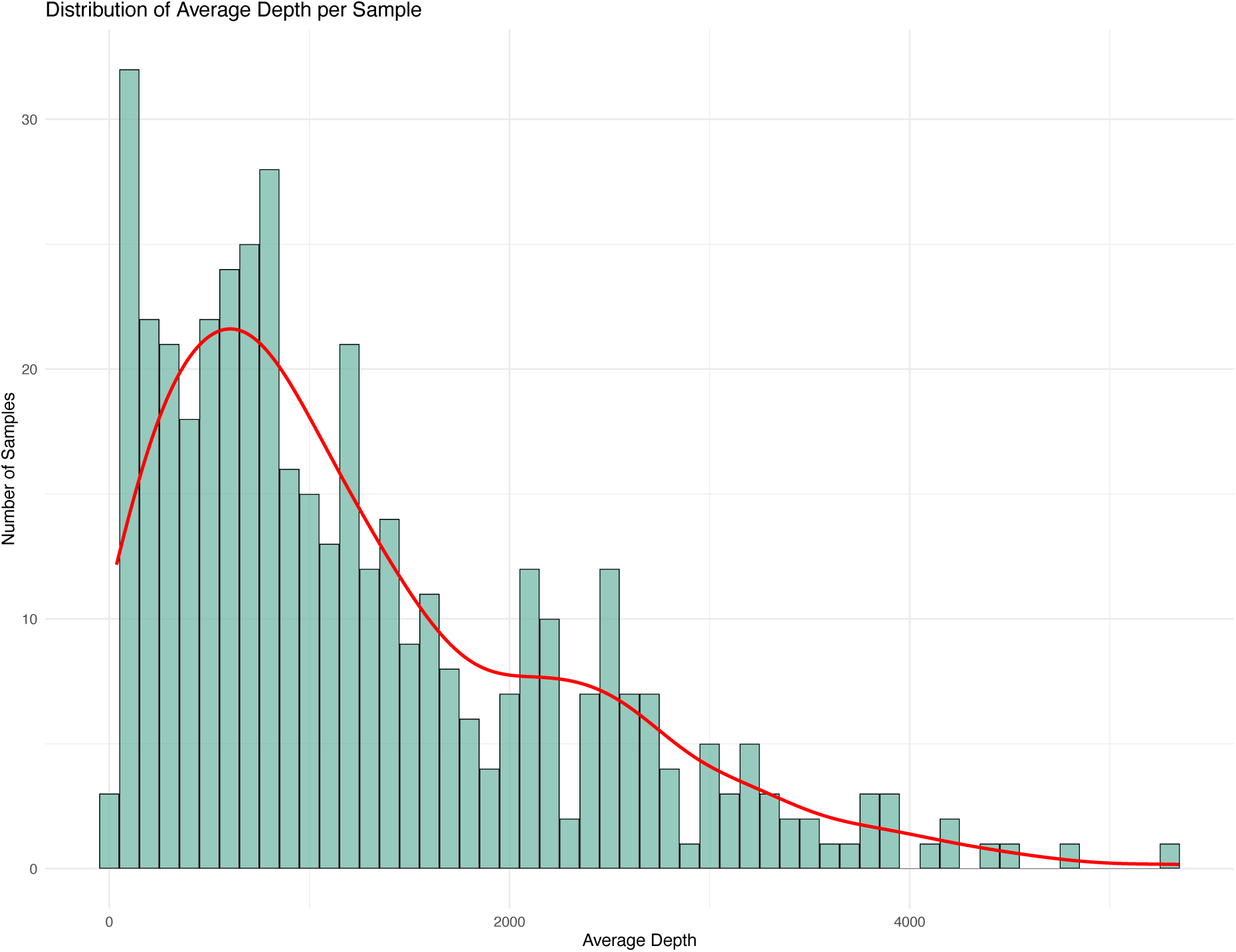
Distribution of read depth across the mitochondrial genome based on whole-genome resequencing data from 430 *Philaenus spumarius* individuals, mapped to the mitochondrial assembly of Phspu_JIC_v2. The x-axis represents the average read depth values, while the y-axis shows the number of individuals exhibiting each read depth.

**Figure S10.**
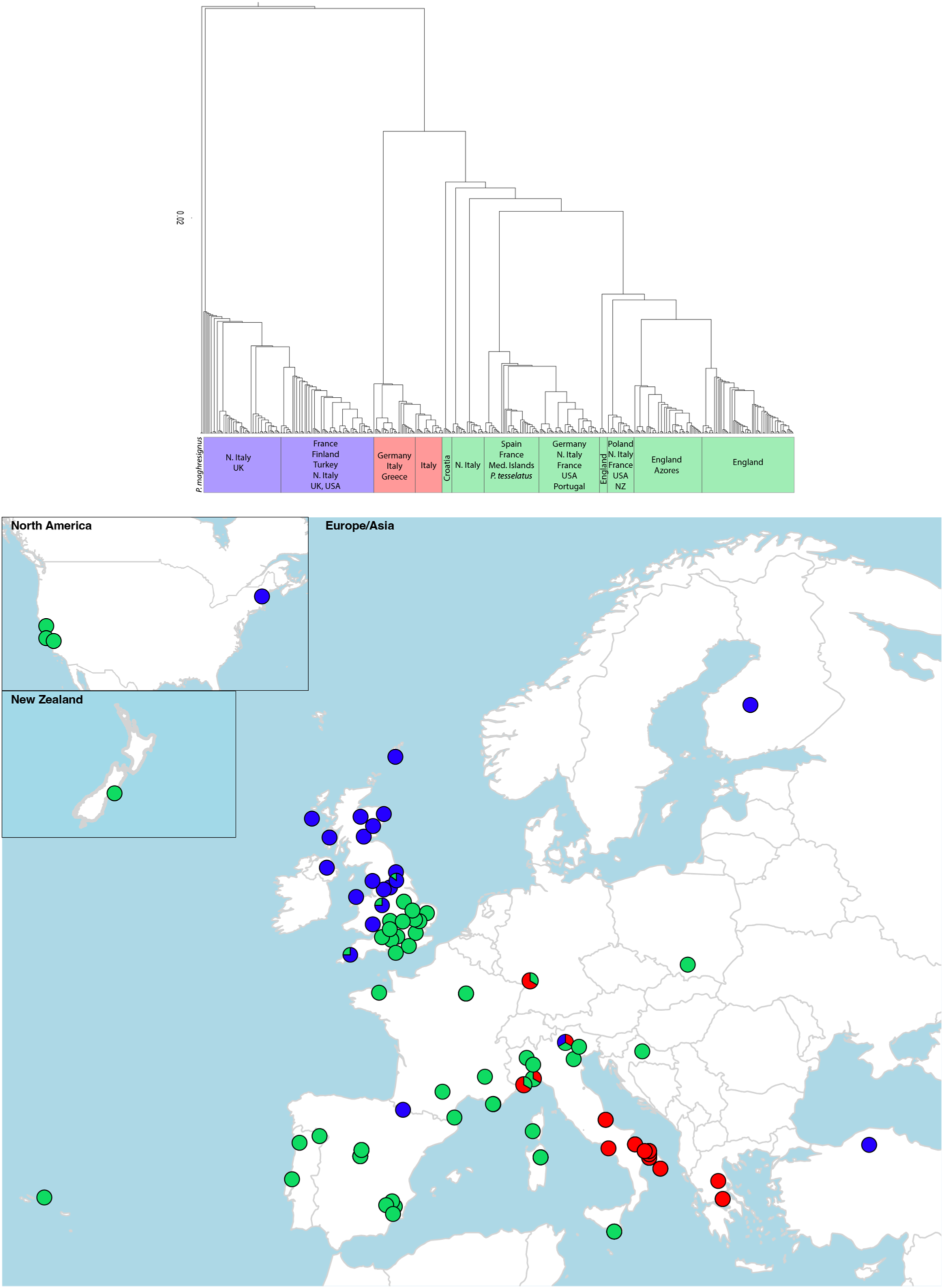
Population structure of 430 *Philaenus spumarius* individuals based on mitochondrial haplogroups. Upper panel, a phylogenetic tree based on mitochondrial genome sequences of 430 *P. spumarius* and one *P. tesselatus* individuals, using that of *P. maghresignus* as the outgroup**. Lower panel**, mitochondrial haplogroup distribution map. Coloured circles denote three primary lineages corresponding to the previously defined Eastern Mediterranean (red), Western (green), and North-Eastern (blue) haplogroups (haplogroups colours follow Rodrigues et al., 2014).

**Figure S11.**
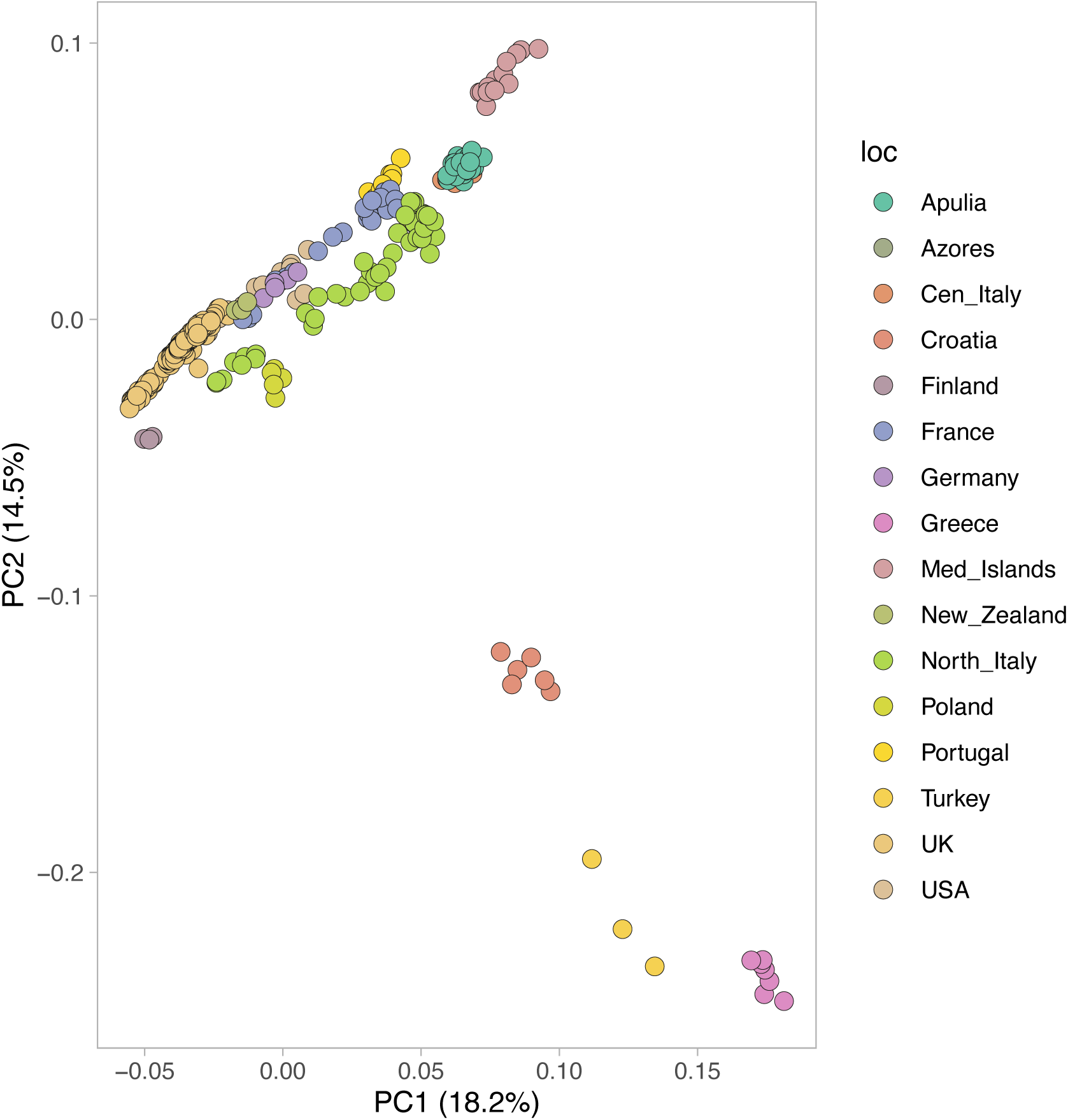
Principal component analysis (PCA) based on genome resequencing data from 409 *Philaenus spumarius* individuals, excluding those sampled in Spain. The plot shows genetic variation along the first two principal components (PC1 and PC2). Individuals from different sampling locations are coloured according to their geographic origin.

**Figure S12.**
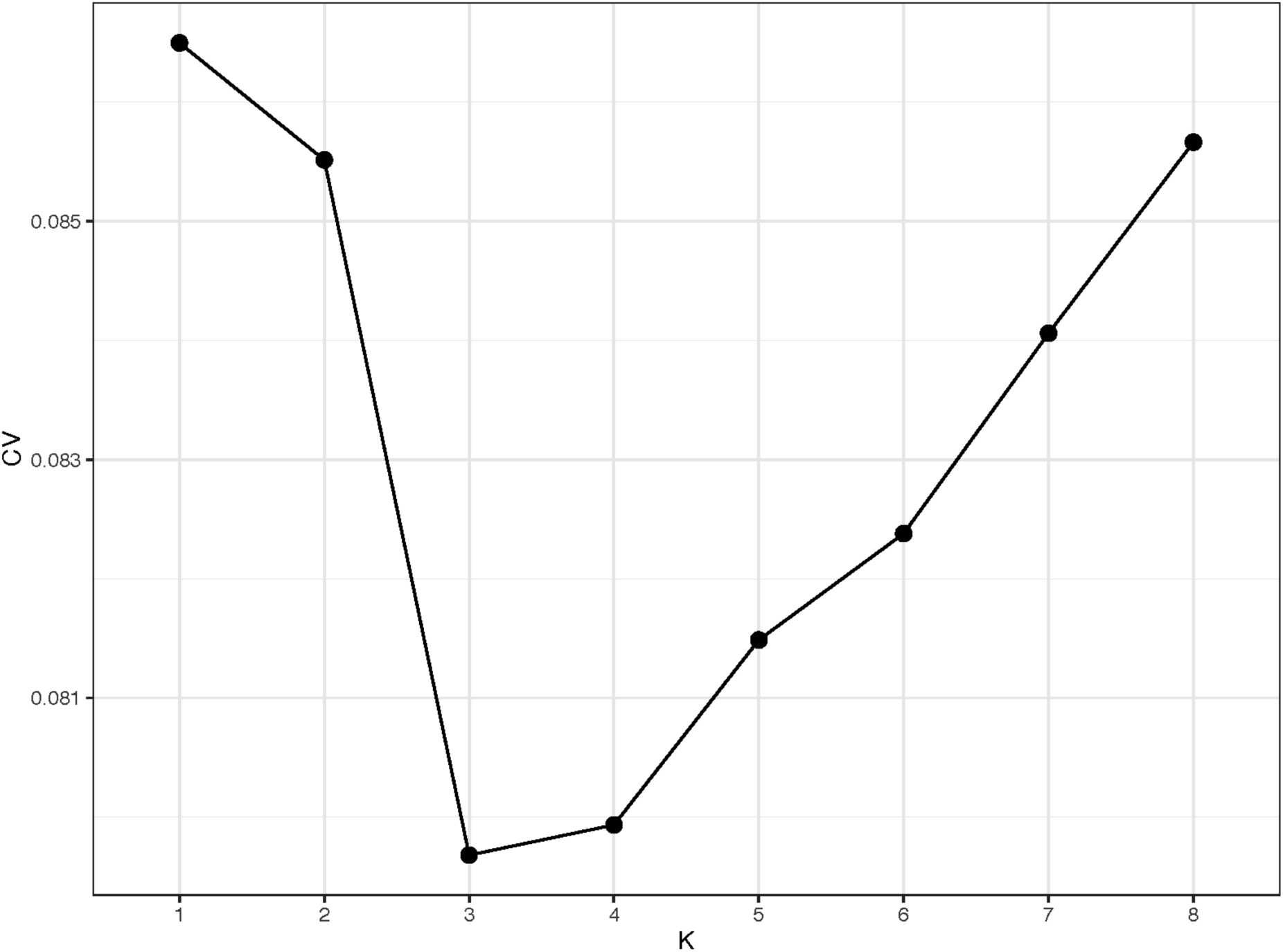
Cross-Validation error rates for admixture runs (K1-K8) based on genome resequencing data from 430 *Philaenus spumarius* individuals. The x-axis represents the tested values of K (number of genetic clusters), and the y-axis shows the corresponding CV error values.

**Figure S13.**
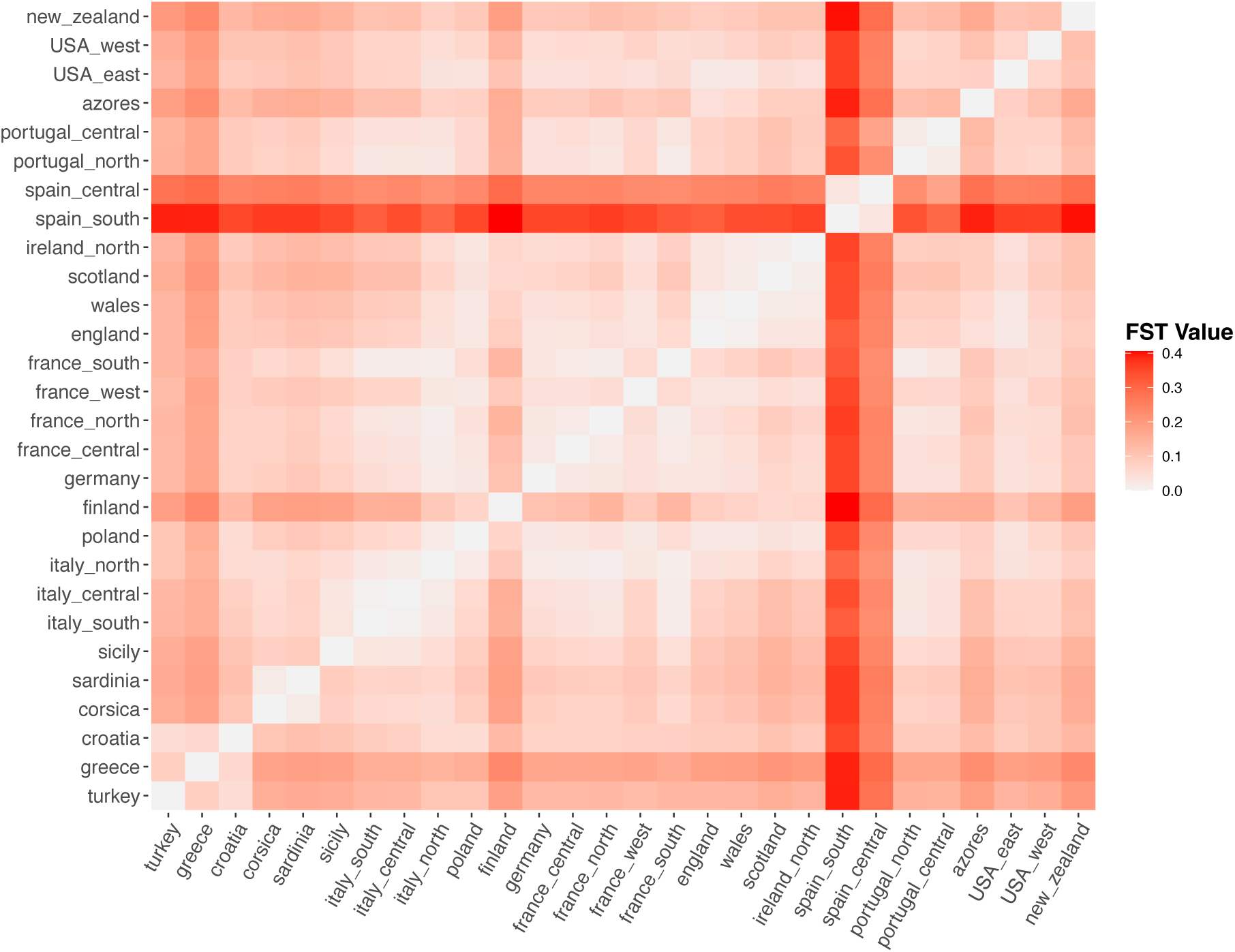
Pairwise F_ST_ comparisons based on whole-genome sequence data of *Philaenus spumarius* collected at main geographical areas. F_ST_ values, calculated in 50-kb windows across the genome, range from 0 (white) to 1 (dark red).

**Figure S14.**
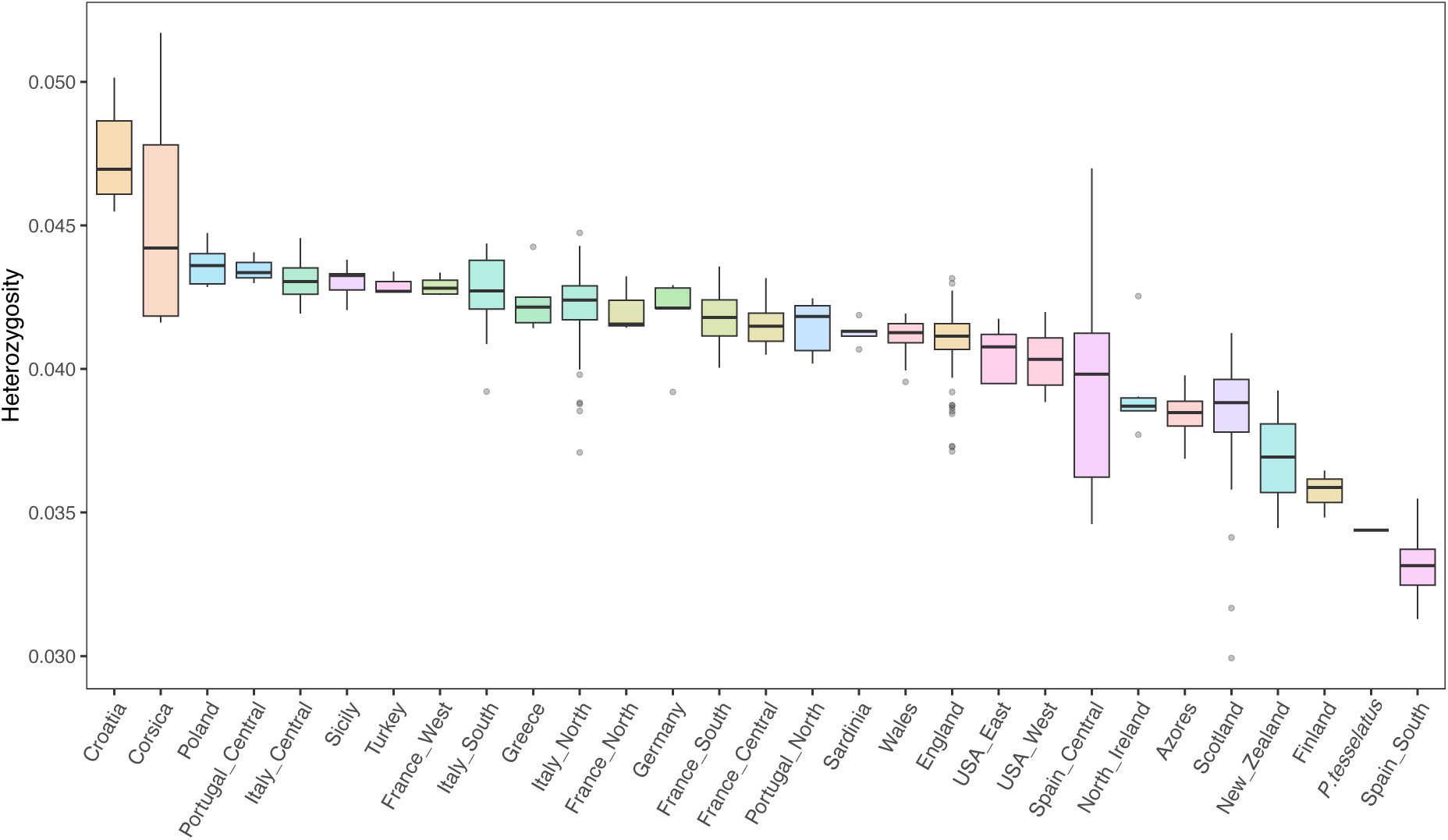
Heterozygosity levels based on whole-genome sequence data of *Philaenus spumarius* and *P. tesselatus* individuals within main geographical areas. The x-axis represents the sampling locations, while the y-axis shows the heterozygosity levels for each main area.

**Figure S15.**
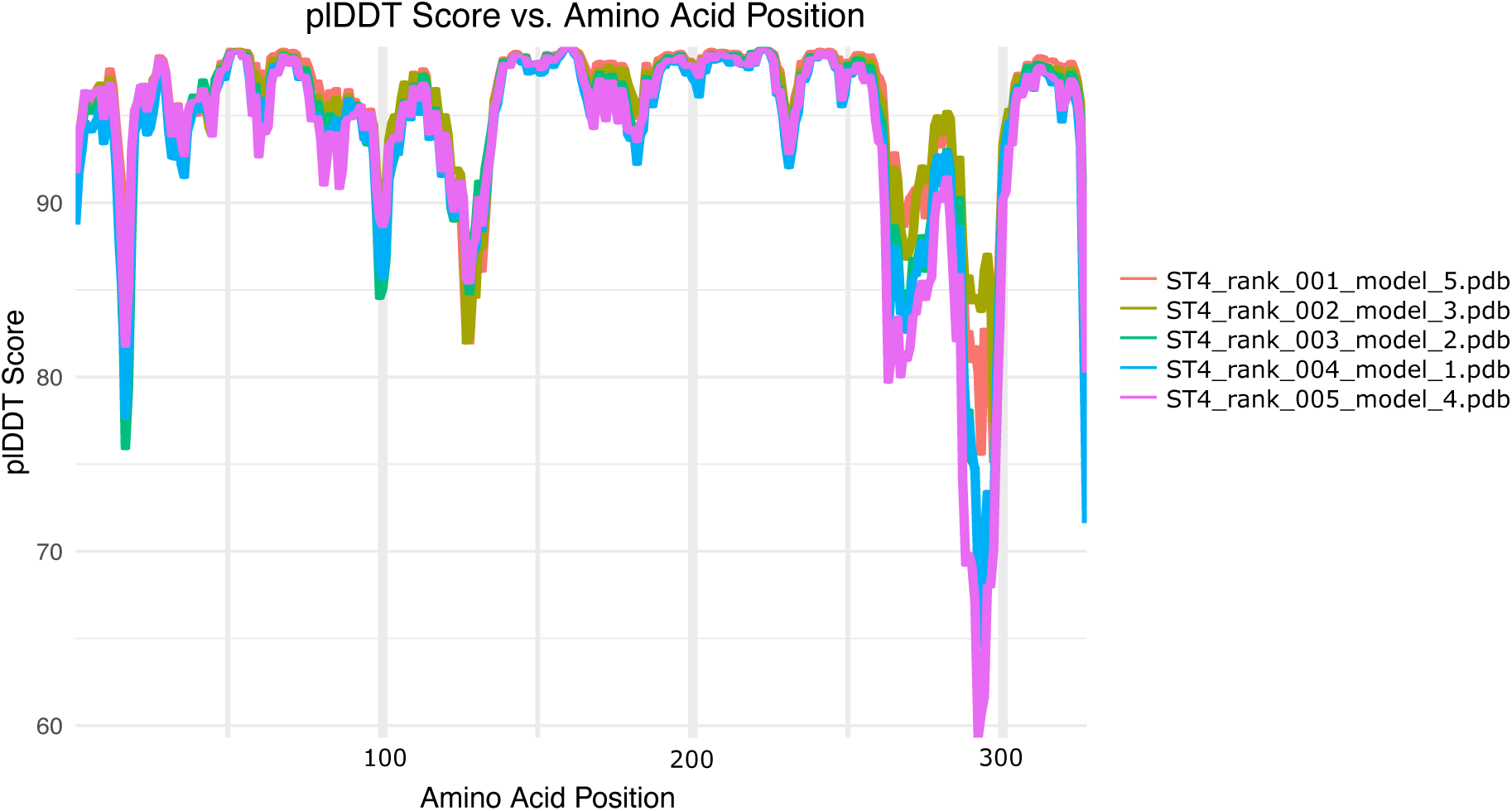
PLDDT scoring model of the ST4 protein structure modelled using Alphafold2. The x-axis represents the amino acid position along the protein sequence, while the y-axis shows the PLDDT score.

**Table S1.** Genome annotation statistics of *Philaenus spumarius, P. tesselatus, P. maghresignus, Aphrophora alni, Cercopis vulnerata and Callitettix versicolor*.

**Table S2.** Transposable element annotations of *Philaenus spumarius, P. tesselatus, P. maghresignus, Aphrophora alni, Cercopis vulnerata and Callitettix versicolor*.

**Table S3.** Insect genomes used for phylogenomic analysis. For each species the taxon ID used in the orthofinder analysis is given.

**Table S4.** Overall summary statistics for OrthoFinder gene family clustering of proteomes for Auchenorrhyncha species listed in Table S3.

**Table S5.** Summary statistics for OrthoFinder gene family clustering of proteomes for each of the Auchenorrhyncha species listed in Table S3.

**Table S6.** Gene duplication numbers in Auchenorrhyncha species listed in Table S3 Table S7. Metadata of insects used for genome sequencing.

**Table S8.** Metadata of insects used for genome resequencing in *P. spumarius* population structure analyses.

